# Tracking clonal evolution of drug resistance in ovarian cancer patients by exploiting structural variants in cfDNA

**DOI:** 10.1101/2024.08.21.609031

**Authors:** Marc J. Williams, Ignacio Vázquez-García, Grittney Tam, Michelle Wu, Nancy Varice, Eliyahu Havasov, Hongyu Shi, Gryte Satas, Hannah J. Lees, Jake June-Koo Lee, Matthew A. Myers, Matthew Zatzman, Nicole Rusk, Emily Ali, Ronak H Shah, Michael F. Berger, Neeman Mohibullah, Yulia Lakhman, Dennis S. Chi, Nadeem R. Abu-Rustum, Carol Aghajanian, Andrew McPherson, Dmitriy Zamarin, Brian Loomis, Britta Weigelt, Claire F. Friedman, Sohrab P. Shah

## Abstract

Drug resistance is the major cause of therapeutic failure in high-grade serous ovarian cancer (HGSOC). Yet, the mechanisms by which tumors evolve to drug resistant states remains largely unknown. To address this, we aimed to exploit clone-specific genomic structural variations by combining scaled single-cell whole genome sequencing with longitudinally collected cell-free DNA (cfDNA), enabling clonal tracking before, during and after treatment. We developed a cfDNA hybrid capture, deep sequencing approach based on leveraging clone-specific structural variants as endogenous barcodes, with orders of magnitude lower error rates than single nucleotide variants in ctDNA (circulating tumor DNA) detection, demonstrated on 19 patients at baseline. We then applied this to monitor and model clonal evolution over several years in ten HGSOC patients treated with systemic therapy from diagnosis through recurrence. We found drug resistance to be polyclonal in most cases, but frequently dominated by a single high-fitness and expanding clone, reducing clonal diversity in the relapsed disease state in most patients. Drug-resistant clones frequently displayed notable genomic features, including high-level amplifications of oncogenes such as *CCNE1*, *RAB25*, *NOTCH3*, and *ERBB2*. Using a population genetics Wright-Fisher model, we found evolutionary trajectories of these features were consistent with drug-induced positive selection. In select cases, these alterations impacted selection of secondary lines of therapy with positive patient outcomes. For cases with matched single-cell RNA sequencing data, pre-existing and genomically encoded phenotypic states such as upregulation of EMT and VEGF were linked to drug resistance. Together, our findings indicate that drug resistant states in HGSOC pre-exist at diagnosis and lead to dramatic clonal expansions that alter clonal composition at the time of relapse. We suggest that combining tumor single cell sequencing with cfDNA enables clonal tracking in patients and harbors potential for evolution-informed adaptive treatment decisions.

## INTRODUCTION

For women diagnosed with advanced high-grade serous ovarian cancer (HGSOC), the prognosis is poor; only 17% will remain long-term disease free^1^ after upfront treatment with platinum-based chemotherapy. Based on the high relative mortality to incidence rate, ovarian cancer ranks as the sixth most lethal malignancy affecting women^2^. Its lethality has been attributed largely to advanced stage at diagnosis, due in part to the absence of effective screening for early-stage disease. Front-line treatment includes surgical resection and combination platinum-taxane chemotherapy, which are initially effective. Nevertheless, most patients will experience recurrence and ultimately die from the disease. Treatment failure in cancer patients is often driven by cancer evolution, owing to selection and expansion of subsets of cells that acquire drug resistant phenotypes^3^. We posit that real-time tracking of cancer evolution in patients has the potential to steer clinical decisions, optimize treatment approaches and discover drivers of drug resistance. Indeed, next generation clinical trial designs are being proposed to investigate how to optimally overcome drug resistance driven by cancer evolution^4^. However, the methods required to monitor evolutionary dynamics in the clinical context are currently lacking. Serial tumor sampling for genomic profiling from multiple time points is often impractical or contra-indicated, making tissue-based longitudinal studies both logistically and clinically challenging. Meanwhile, powerful techniques like cellular barcoding provide insights in model systems^5–7^ but cannot be applied in patients. Non-invasive serial imaging and blood-derived biomarkers provide other sources of longitudinal information, but these lack tumor cell-intrinsic molecular measures needed to capture the intra-tumor heterogeneity for monitoring evolutionary dynamics. Recent advances in cell-free DNA (cfDNA) profiling to detect tumor-derived DNA from routinely collected blood samples has changed the field of non-invasive molecular diagnostics for cancer patients^8–10^. Here, we demonstrate that tracking evolutionary dynamics in cfDNA from HGSOC diagnosis to recurrence can be implemented in patients as a powerful evolution-centred tool to study the molecular determinants of drug resistant relapsed disease *in vivo*.

Using cfDNA to study cancer evolution in patients is a relatively nascent field^11,12^. The main objective is to first identify clonal populations, and subsequently use their clone-specific genotypes as endogenous markers to estimate the relative tumor fraction of each clone in cfDNA over time. Here, we contend that tumor tissue sequencing as the basis of identification of clonal populations which can then be tracked over time by sequencing serially collected cfDNA samples is a route to precisely monitor disease evolution. In contrast to clonal decomposition from bulk sequencing, which is imprecise^13^ especially when tumor content and/or sequence coverage is low, single cell whole genome sequencing (scWGS) approaches have shown great promise in unambiguously resolving clonal composition^14,15^. In particular, shallow whole genome sequencing technologies provide reliable readouts of clonal composition^16–18^, especially in cancer types such as HGSOC characterized by genomic instability^17,18^, even resolving clones to approximately 1% prevalence^19–21^. Furthermore, by combining clonally related cells into pseudobulk, point mutations and structural variant breakpoints can be identified, providing clone-specific genomic features at base-pair resolution^17,18^ which can serve as endogenous barcodes for tracking clonal abundance over time.

Here we show that scWGS on tumor tissue combined with cfDNA clonal tracking is a powerful approach to reveal insights into drug resistance. Our results indicate that i) clonal tracking exploiting clonal structural variations is tractable for monitoring HGSOC disease evolution in patients; ii) drug resistance at relapse is consistent with clonal pruning and reduced clonal diversity; iii) positive selection operates in the majority of patients leading to near clonal sweeps of high fitness clones; iv) positively selected clones harbor clone-specific high level amplifications of oncogenes including *ERBB2*, *RAB25*, *CCNE1*, *NOTCH3* and a *BRCA1* reversion mutation. Together, these results establish single cell-informed clonal tracking in cfDNA as a powerful approach to measuring and modeling the evolutionary dynamics of relapsed disease in HGSOC, and implicate rare, but pre-existing clones with oncogene amplifications as a putative pre-adapted reservoir of drug resistant cellular populations.

## RESULTS

### Cohort and data generation

We carried out a multi-modal prospective study as part of the MSK SPECTRUM cohort^22,23^, involving 19 newly diagnosed, treatment-naive patients with FIGO stage III/IV HGSOC, with diagnosis verified through clinicopathological review. Patients were followed over a period of up to 5 years and plasma cfDNA was collected during treatment and at the time of radiologic disease recurrence. All 19 patients had cfDNA collected at or close to the time of first debulking surgery or laparoscopic biopsy (baseline), and a subset (n=10) had radiographically confirmed disease recurrence along with cfDNA collections post-recurrence and during therapy (**Supplementary Figure 1**). At the time of tissue collection, fresh tissue samples were collected from multiple disease sites from primary debulking surgeries for patients receiving adjuvant chemotherapy and from laparoscopic biopsies taken at diagnosis for patients undergoing neoadjuvant chemotherapy. Tissues were processed for scWGS with the DLP+ protocol^17^. See **Supplementary Figure 1** for clinical details, treatment history and sample collections for all 19 patients.

### Clone-specific mutations and structural variations in scWGS

From the 19 patients included in this study we generated scWGS data from 19,454 cells (range 200-2015 cells per patient, **Supplementary Figure 2**) with mean coverage of 0.089X (range 0.002-0.392X per cell, **Supplementary Figure 2**). We inferred the clonal composition at the time of diagnosis based on copy number data (**Methods**), with the aim of following these clones over time as patients received chemotherapy, maintenance therapies and experienced disease recurrences using cfDNA (**Supplementary Figure 3**).

To follow clones over time in cfDNA we identified clone-specific markers, structural variants(SVs) and single nucleotide variants(SNVs) in each patient. Due to the sparse coverage in scWGS data, the presence/absence of SNVs and SVs cannot be determined in every cell. We therefore developed a combination of pseudo-bulk mutation calling and single-cell copy number phylogenetics to confidently identify clone-specific mutations that could be profiled in cfDNA, focusing primarily on SVs resulting from genomic rearrangements. As SVs are a hallmark of HGSOC genomes, we reasoned they would provide a highly specific readout in cfDNA due to their unique sequence composition, where breakends juxtapose sequence from distal chromosomal loci. As a result, these unique sequences should be largely immune to sequencing error and other causes of false positive detection in cfDNA.

To begin identifying clone-specific SVs, we first constructed single-cell phylogenies with `MEDICC2`^24^ using allele-specific copy number alterations as input (500kb resolution, see **Fig. 1a** for patient 004). Clones were defined based on divergent clades from the single-cell phylogenetic trees (**Methods**). We then merged cells from each clone and re-computed copy number at 10kb resolution using a new Hidden Markov Model (HMM) based copy number caller, `HMMclone` (**Methods**). `HMMclone` improves the resolution of pseudobulk clone copy number profiles and enables more precise matching between copy number and SVs (**Supplementary Figure 4**). SVs and SNVs were identified in sample-level ‘pseudobulk’ data and genotyped in single-cells (**Methods**). Although only a small proportion of cells (<5%) have reads that support a mutation or SV of interest, we tested whether the distribution of the subset of cells positive for a mutation across clones in the tree could inform mutation clonality. For example, a truncal missense *TP53* mutation and a truncal 1.03Mb deletion in 004 distributed uniformly across the tree and were present across all clones (**Fig. 1b,c**). Cells with support for subclonal clone-specific mutations on the other hand – in this case 2 SNVs and 2 duplications – distributed non-randomly in a clone-specific manner (**Fig. 1b,c**). This ‘parsimony’ principle extended to more complex events, for example a chromothriptic-like chr8 in this patient. Clone-specific pseudobulk copy number at 10kb resolution showed that the chromothripsis, although sharing some common features, is divergent between clone A and clone B (**Fig. 1d**), providing a rich source of SVs that are clone-specific.

**Figure 1.**
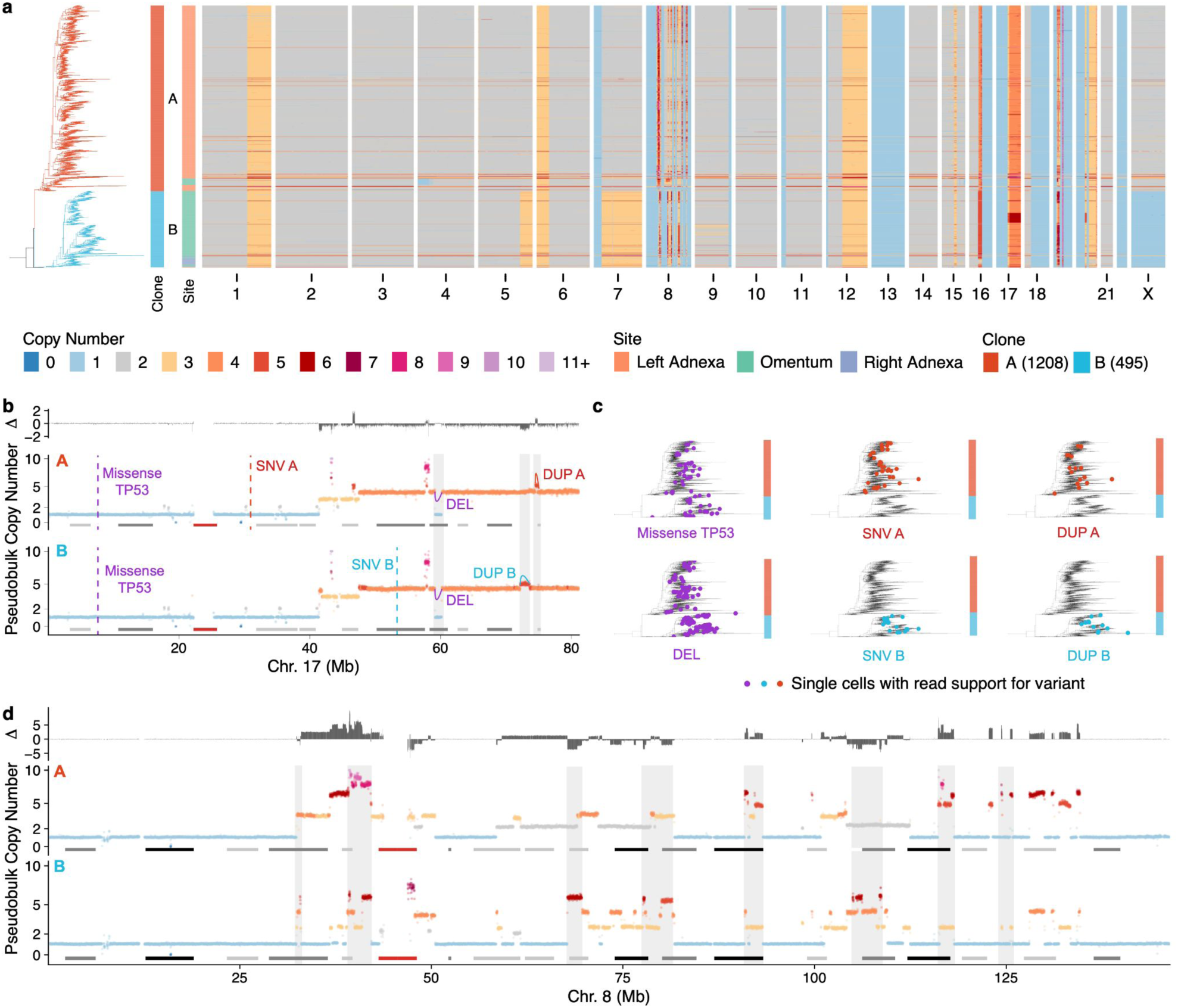
Clone-specific mutations and structural variations in scWGS. **a)** scWGS based copy number heatmap for patient OV-004. Each row is the copy number of a cell, cells are ordered according to a MEDICC2 computed single-cell phylogeny (shown on the left) **b)** Clone pseudobulk copy number at 10kb resolution for clone A and clone B in chr17. Truncal variants (*TP53* missense and deletion) are annotated in purple, clone specific duplications and SNVs are annotated in red and blue respectively **c)** Phylogenetic trees annotated with cells that have support for variants shown in panel **b)**. **d)** Clone pseudobulk copy number at 10kb resolution for clone A and clone B in chr8 showing different chromothriptic chromosomes. In **b)** and **d)** notable regions that are different between clones A and B are highlighted in gray.

### Structural variants as highly specific markers of tumor DNA in cfDNA

With clone-specific SVs identified, we then determined the utility of SVs as markers of tumor DNA in plasma, and compared their quality and robustness relative to SNVs that have been the focus of most cfDNA assays, including commercial ones. For each patient, we constructed a panel of mutations comprising a mix of clonal and subclonal somatic SNVs/SVs (≥100 SVs per patient, **Supplementary Figure 2**) and a small number of germline single nucleotide polymorphisms (SNPs) for QC purposes. We designed patient-bespoke hybrid capture probes with 60bp flanking sequence on either side of the breakpoint or point mutation, and incorporated these probes into a cfDNA duplex error-corrected sequencing assay^25^ (mean raw coverage 13,531X; mean consensus duplex coverage 970X, **Fig. 2a**). To estimate baseline accuracy, we first applied the assay to cfDNA plasma samples taken at or close to the time of tissue collection, assuming tumor burden and thus tumor-derived cfDNA yield would be high. For benchmarking purposes, we characterized the sensitivity and error profiles of truncal mutations that were also detected in matched bulk whole genome sequencing data. For example, reads supporting a truncal translocation between chr8 and chr19 in patient 107, were easily identified as they aligned across the breakpoint in cfDNA, single cells and bulk tumor whole genome sequencing (**Fig. 2a**). Across all 17 pre-operative baseline cfDNA samples with sufficient SNVs for comparison, ctDNA with SNVs and SVs were detected and VAF distributions derived from the error corrected sequences were concordant between SNVs and SVs (**Fig. 2b**).

**Figure 2.**
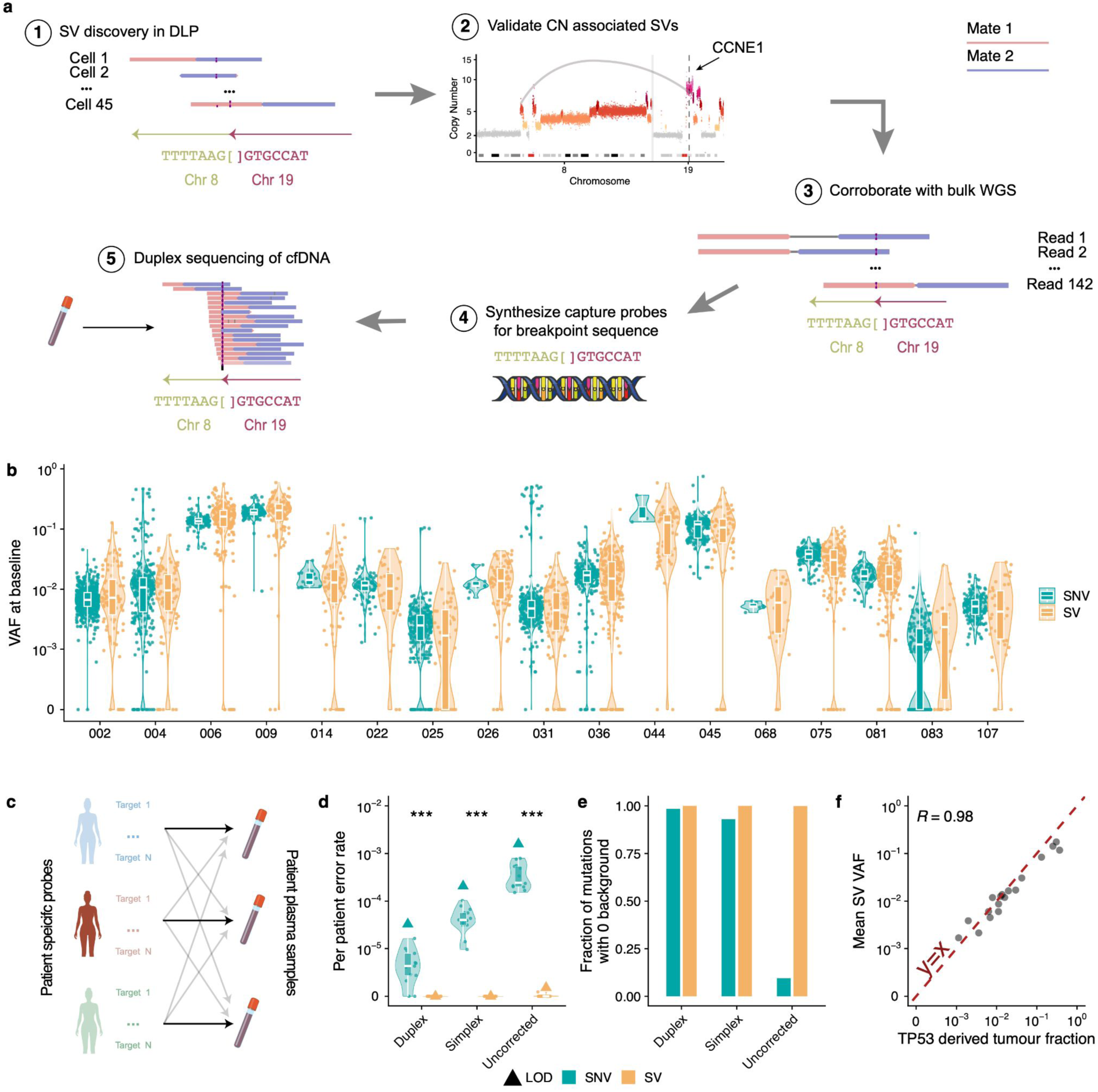
Structural variants as highly specific markers of tumor DNA in cfDNA. **a)** Schematic of workflow illustrated with a translocation between chr8 and chr19 identified in OV-107. **b)** Distribution of VAFs for SVs and SNVs in baseline samples **c)** Schematic showing how patient specific error rates are calculated by applying probe sets to off target patients **d)** average background error rates in duplex, simplex and uncollapsed sequences. Each violin/boxplot is a distribution over SVs/SNVs where each data point is the error rate for an individual patient. Triangles show limit of detection (LOD) defined as 2X the largest observed patient error rate **e)** Fraction of SNV/SVs that have 0 background ie no read support in incorrect patient **f)** Mean SV VAF vs Tumor fraction computed from TP53 VAF

To compute background error rates, patient-specific hybrid capture probe sets were applied to the ‘on-target’ patient as well as at least one other ‘off-target’ patient (**Fig 2c**), where we expect no detection. Background error rates were defined as the total number of off-target variant supporting reads divided by the total number of reference reads per patient. Error rates were computed for duplex sequences (collapsing reads from both strands of the initial cfDNA molecule), simplex sequences (one strand) and the raw uncorrected sequences. Background error rates were negligible for SVs; we observed no errors in duplex or simplex sequences (**Fig. 2d**). In the uncorrected sequences we observed read support for a single event from one patient (**Fig. 2d**). Compared to SNVs whose error rates increased in simplex and uncorrected sequences relative to duplex sequences as expected, error rates for SVs were orders of magnitude lower and were negligible even in uncorrected sequencing (**Fig. 2d,e**, p<10^-10^, t-test). Using this data we defined the limit of detection (LOD) as 2X the largest observed patient error rate (**Fig. 2d)**. Given that we observed no errors for duplex or simplex sequences, we determined the upper bound for the LOD to be ∼10^-7^ (inverse of the total number of reference supporting reads). The mean VAF of SVs was correlated with tumor fraction (R=0.98, p<10^-10^, pearson correlation) calculated from *TP53* mutation VAF measurements, assumed to be truncal in all HGSOC patients^26^ (**Fig. 2f**). Together, these data demonstrate that SVs can be readily detected in plasma cfDNA, have a lower background error rate compared to SNVs and can be used to estimate tumor fractions.

### Detecting clone-specific SVs in cfDNA

We next tested whether clone-specific SVs can be detected in plasma cfDNA. Clone-specific SVs inferred by scWGS analysis were present in all patients with at least 200 cells (18 patients, average n=144, range 29-361 **Supplementary Figure 2**). We found that numerous mutational processes such as chromothripsis^27^ (e.g. patient 083, **Fig. 3a**), breakage fusion bridge^28^-induced focal amplifications (patient 045, **Fig. 3b**), pyrgo-like tandem duplication “towers”^29^ (consequence of *CDK12* mutant tandem duplication phenotype; patient 081, **Fig. 3c**) and complex intra-chromosomal^30^ events (patient 002, **Fig. 3d**) contributed to clone-clone differences in SVs. Clone-specific SVs were co-located with copy number changes as expected (**Fig. 3a-d**). Using the probe designs as described above, clone-specific SVs were detected in all baseline plasma cfDNA samples (**Fig. 3a-d**), even in samples with tumor DNA fractions <1% (**Fig. 3e,f**) and VAFs of subclonal variants were lower relative to clonal variants as expected (p < 0.001 in 16/18 patients, n.s. in 2/18, t-test, **Fig. 3f**), supporting the clonal structure found in the tissues. These results therefore establish that scWGS enables accurate assignment of SVs to clones, that SVs are sensitive markers of tumor DNA and that clone-specific SVs can be detected in low tumor fraction plasma.

**Figure 3.**
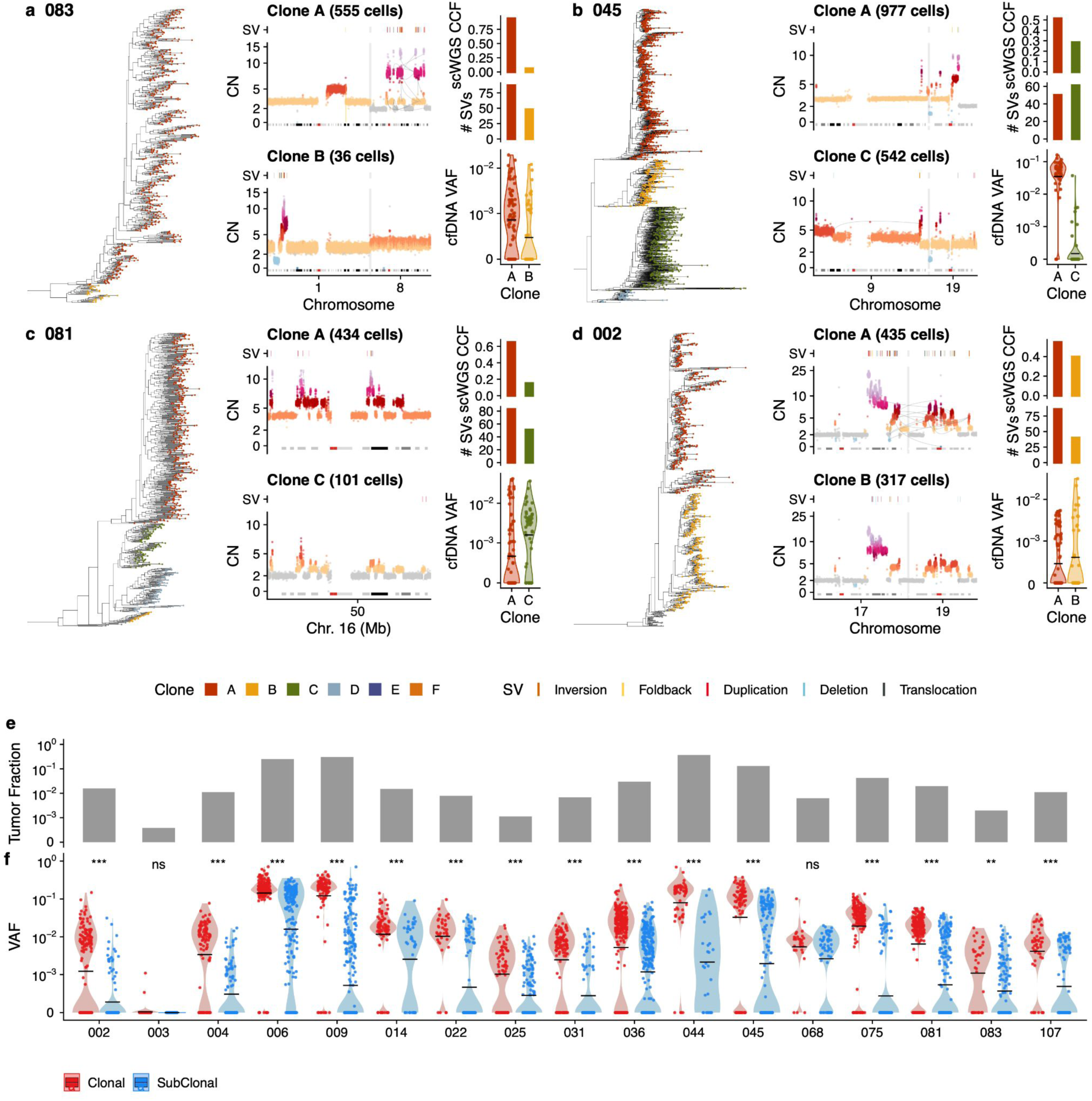
Detecting clone-specific SVs in cfDNA. a)-d) Single cell phylogeny on left hand side with tips coloured by clone membership, zoom in on copy number profiles of chromosomes of interest that have clone specific structural variants driven by a mutational process, above each copy number profile, the location of SVs are shown, right hand side shows the CCF of the 2 clones of interest in DLP, the number of clone specific structural variants and the VAF of those clone specific SVs in cfDNA at baseline. Shown are chromothripsis in OV-083, breakage-fusion bridge in OV-045, tandem duplication towers in OV-081 and chromoplexy in OV-002. e) Tumor fraction in baseline samples inferred from TP53 mutation f) VAF of all structural variants at baseline in cfDNA stratified by clonality. Black horizontal line shows mean value.

### Clonal evolution of drug resistance in patients

We next evaluated whether our approach could be used for longitudinal monitoring of tumor evolution during treatment and disease recurrence. From our patient cohort, we studied 10 patients with radiographically confirmed disease recurrence and profiled all available post-baseline and post-recurrence cfDNA plasma samples with our patient-specific assay (mean 7.8 timepoints per patient, range 3-13). See **Supplementary Figure 5** for the scWGS data for these 10 patients. For all patients, ctDNA VAF of truncal SVs decreased during initial chemotherapy, as patients responded to therapy with decreased burden and decreased serum CA-125 levels (**Fig. 4** & **Supplementary Figure 6**). All patients were positive for ctDNA at the time point closest to first recurrence (defined as average VAF across the panel exceeding LOD (**Fig. 4** & **Supplementary Figure 6**)). In 6 patients with sufficient plasma samples, ctDNA was detected prior to clinically confirmed disease recurrence but subsequent to completion of initial chemotherapy: 002 (76 days), 004 (26 days), 009 (184 days), 045 (109 days), 075 (233 days), 081 (314 days) (**Fig. 4** & **Supplementary Figure 6**). We note that not all of these patients achieved ctDNA clearance (045, 075, 081), which may in part be due to insufficient cfDNA sampling at completion of first-line chemotherapy.

**Figure 4.**
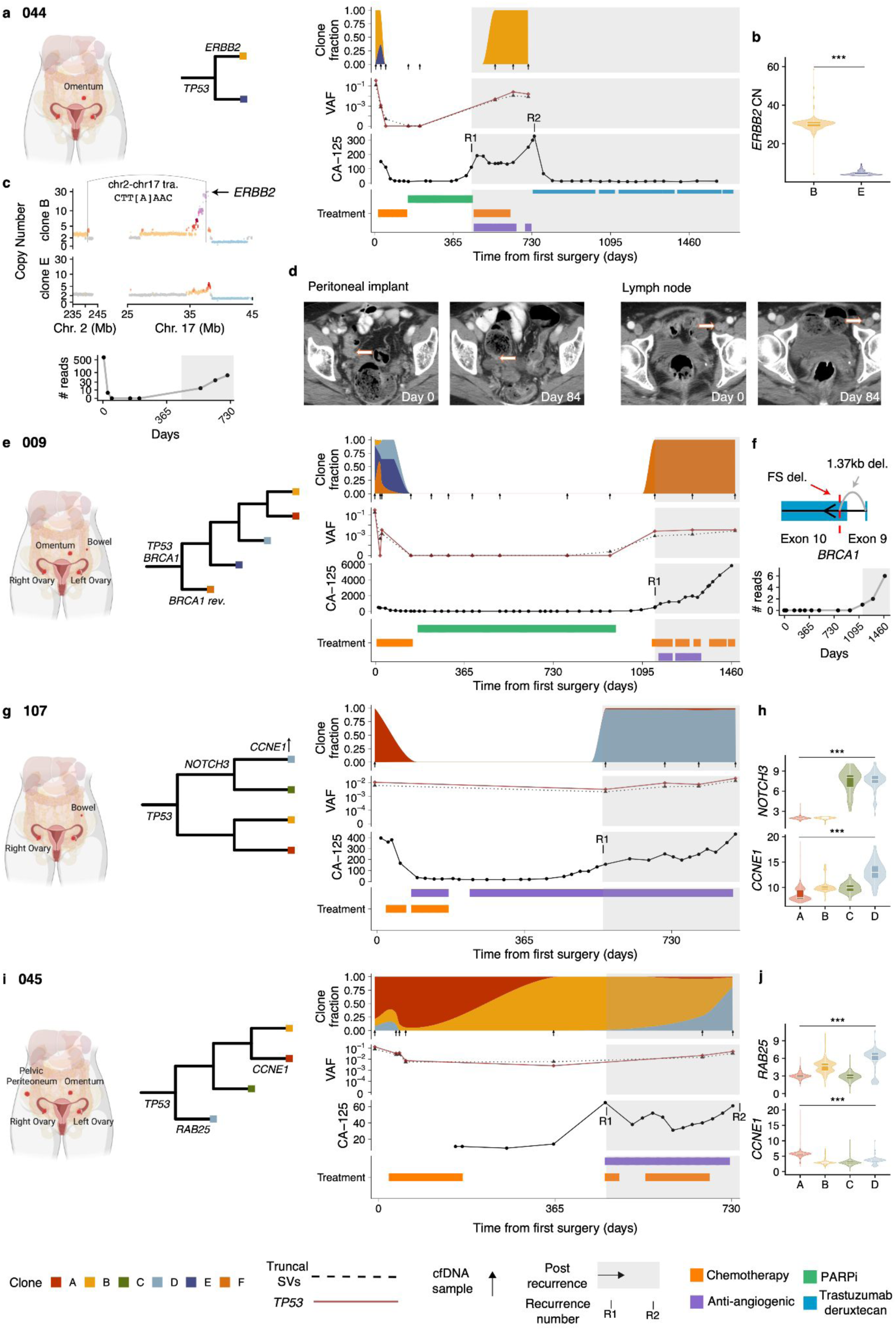
Clonal evolution of drug resistance in patients. Clonal evolution tracking in 4 patients. a) Anatomical sites sequenced with DLP, a phylogenetic tree of the clones, then clonal fractions, mean truncal SV VAF and *TP53* VAF, CA-125 and treatment history over time for patient 044. Disease recurrences are annotated on the CA-125 track. b) *ERBB2* copy number in clone B vs E across cells c) Pseudobulk copy number of clones B and E at 10kb resolution in chromosomes 2 and 17. A translocation specific to clone E and implicated in the ERBB2 amplification is highlighted. Below shows the read counts of this translocation across timepoints in cfDNA d) CT scan images from day 0 and day 84 from 2 sites. Orange/white arrows indicate site of disease e) Clonal tracking in patient 009, same as panel a). f) Diagram of mutations impacting the *BRCA1* gene: location of frameshift deletion shown with red dashed line, large 1.37kb deletion shown in gray. Number of reads supporting the 1.37kb deletion in cfDNA across time. g) Clonal tracking in patient 107, same as panel a). h) *NOTCH3* and *CCNE1* single cell copy number distribution across clones i) Clonal tracking in patient 045, same as panel a). j) *RAB25* and *CCNE1* single cell copy number distribution across clones

We then measured how the abundance of specific clones changed over time as patients received treatment. Clone abundances at each time point were estimated by averaging the VAF across all structural variants assigned to a clone. Firstly, to validate the accuracy of inferred clone frequencies we performed WGS of plasma at 20X coverage from 6 samples (1 baseline and 1 recurrence sample from 3 patients: 045, 081 and 107). Copy number profiles from these data were consistent with predictions derived from scWGS derived copy number profiles and ctDNA clone frequencies (**Supplementary Figure 7a-c).** Notably, clone-specific amplifications, which provide the strongest signals in such low tumor fraction sequencing data, were consistent with the inferred dominant clone at baseline and recurrence. In patient 045, the *CCNE1* locus (chr19q) was enriched at baseline, and *RAB25* (chr1q) at recurrence (**Supplementary Figure 7d**), while in patient 107, *CCNE1* and a region on chr20q were enriched at recurrence as expected (**Supplementary Figure 7e**). As further validation, we computed clonal frequencies across time using SNVs for patients with sufficient clone-specific SNVs (minimum of 4 per clone) and compared them to the clone frequencies estimated using SVs. Clone frequencies across time for patient 045 showed highly similar patterns for SVs (**Supplementary Figure 7f**) and SNVs (**Supplementary Figure 7g**) and were consistent using these two distinct sets of genomic features for all patients (**Supplementary Figure 7h**, R=0.93, p<10^-10^, Pearson correlation).

Having confirmed the accuracy of our inferred clonal trajectories we then aligned clone frequencies to treatment histories and other clinical biomarkers such as serum CA-125 levels enabling us to precisely describe clonal evolution in the context of therapy and disease recurrence. In patient 044, 2 major clones were present at the time of diagnosis (clone B and clone E, **Fig. 4a**). From the scWGS data we noted that clone B had an *ERBB2* high-level amplification (∼30 copies) that was absent in clone E (**Fig. 4b,c)**. The patient responded to upfront chemotherapy and achieved ctDNA clearance at day 156 along with a notable drop in CA-125 level. Reduction in disease burden due to upfront chemotherapy can be seen from radiology images of the same disease sites at day 0 vs day 84 (**Fig. 4d)**. The patient then experienced disease recurrence at day 449 and received second line chemotherapy. Post-recurrence cfDNA samples during the second line of chemotherapy only detected clone B (*ERBB2* amplified), and the patient had minimal response to this second line as evidenced by high levels of ctDNA detected and persistently elevated CA-125 (**Fig. 4a)**. These dynamics could be captured by following a single translocation between chromosomes 2 and 17 that was associated with the *ERBB2* amplification (**Fig. 4c)**. Subsequently, following a second disease recurrence at day 730, the patient was treated with trastuzumab deruxtecan, an antibody-drug conjugate that targets HER2 (*ERBB2*). She achieved a complete radiologic response and remains disease free nearly three years after starting therapy. Of note, she was eligible for treatment with trastuzumab deruxtecan on a clinical trial based on clinical tumor-normal MSK-IMPACT^31^ targeted sequencing performed on tissue at the time of diagnosis. As MSK-IMPACT is a bulk assay it would not have identified that there was a clone that lacked the *ERBB2* amplification as was possible with scWGS. The clonal tracking data indicates that front-line therapy eradicated the *ERBB2*-WT clone, leaving the dominant trastuzumab deruxtecan-susceptible *ERBB2*-Amp clone present at recurrence, thus resulting in an exceptional and durable response.

In patient 009, all 5 clones identified in scWGS were detected in cfDNA at diagnosis. The patient experienced a good response to chemotherapy and achieved ctDNA clearance accompanied by a drop in CA-125 levels by day 125 (**Fig. 4e**). The patient had a germline *BRCA1* mutation and received standard of care PARP inhibitor (PARPi) maintenance after completion of chemotherapy, and remained disease free for almost 3 years. ctDNA was detected 184 days prior to clinical recurrence by CT at day 1146. Post recurrence, the only detectable clone was clone F. We identified a putative BRCA1 reversion mutation in post recurrence cfDNA samples; a 1.37kb deletion that excises the beginning of exon 10 (including the germline pathogenic mutation) and the intronic region between exons 9 and 10 of *BRCA1* and restores the reading frame (**Fig. 4e**). We did not find evidence of this event in any of our sequencing data from baseline surgical samples and it was only observed in post-recurrence cfDNA samples (**Fig. 4f**). This event may therefore have been acquired later in a cell from clone F, or alternatively was beyond the limit of detection of our assay at early time points. Of note, patients who experience disease progression following PARPi therapy demonstrate a poor response to subsequent platinum-based chemotherapy^32,33^. Whether this is related to *BRCA1/2* reversion mutations, and how this may impact subsequent clinical care remains an area of active study.

Two patients had clone-specific *CCNE1* amplifications (**Fig. 4g-j**), an alteration that has previously been associated with disease recurrence and chemoresistance in HGSOC^34,35^. In patient 107, clone D had an average *CCNE1* copy number of 13 compared to 8 in clone A (**Fig. 4h**), while in patient 045, *CCNE1* amplification was specific to clone A (6 copies, **Fig. 4j**). In patient 107, clone D was the dominant clone at recurrence, and had a multimegabase amplification on chr19p including *NOTCH3* in addition to increased *CCNE1* copy number on chr19q (**Fig. 4g,h**). Interestingly, in patient 045, although the *CCNE1* amplified clone was dominant at diagnosis, post recurrence and during a second line of chemotherapy, clone D (lacking *CCNE1* amplification) expanded and was the dominant clone at the final time point close to the time of a second disease recurrence (**Fig. 4i**). Although lacking *CCNE1* amplification, clone D harbored an amplification of *RAB25*, a GTPase previously implicated in chemotherapy drug resistance^36^ (**Fig. 4j**). Notably, these results suggest that *CCNE1* amplification at baseline is not deterministically linked to chemotherapeutic resistance.

### Clone-specific transcriptional programs

We next investigated phenotypic associations with drug resistant states, leveraging previously published patient matched scRNAseq data^23^. We first used TreeAlign^37^ to map cancer cells profiled by scRNAseq to genomically defined clones derived from the scWGS data, using all patients in the MSK SPECTRUM cohort for which we had scWGS^22^. For 20 patients for which we could identify at least 2 clones with >100 cells, we then scored each clone by its expression of hallmark pathways and explored how these transcriptional programs varied across clones within the same patient. We found that transcriptional programs could be highly variable between clones from the same patient, suggesting that HGSOC generally has a large degree of pre-existing genomically encoded transcriptional heterogeneity (**Fig. 5a**).

**Figure 5.**
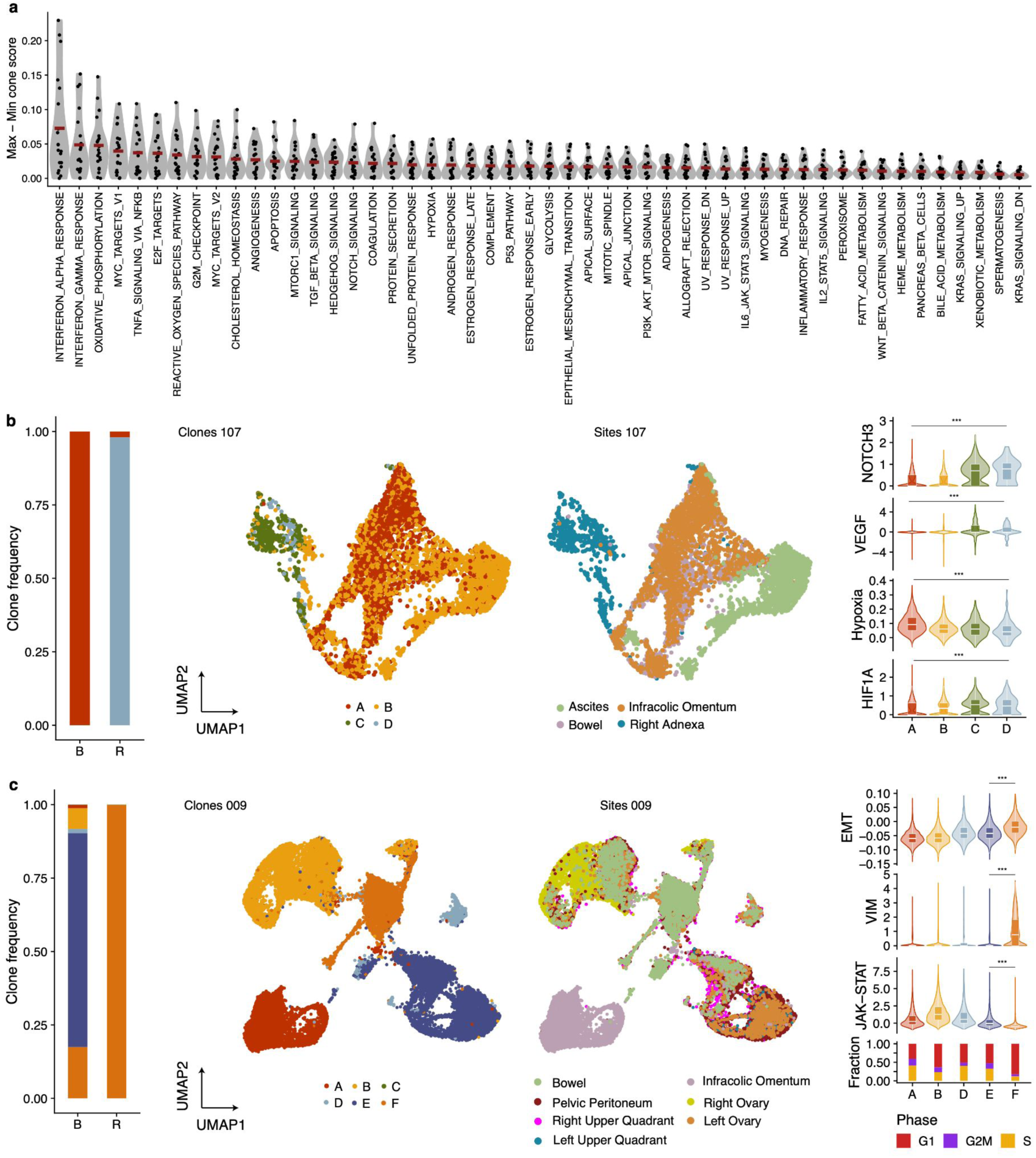
Clone-specific transcriptional programs. **a)** Hallmark pathway variability across genomically defined clones in scRNAseq data. Each data point represent the maximal pathway score difference between clones in each patient. Data from 20 patients included. **b)** From left to right, clone frequencies inferred from cfDNA at baseline (B) and recurrence (R) for OV-107. UMAPs labelled by sites and clone mapping (inferred using TreeAlign). Distribution of NOTCH3 expression, VEGF pathway, hypoxia and HIF1A across clones **c)** Clone frequencies inferred from cfDNA at baseline (B) and recurrence (R) for OV-009 UMAPs labelled by sites and clone mapping (inferred using TreeAlign). Distribution of EMT pathway, VIM expression, JAK-STAT pathway and fraction of cells in each cell cycle phase.

Of the ten patients we had profiled with our longitudinal cfDNA assay, two patients (107 & 009) had sufficient cells assigned to each clone that we could contrast these phenotypic differences in drug resistant versus drug susceptible clones. In patient 107, all 4 clones were represented in the scRNAseq data and clone D was the dominant clone at relapse (**Fig. 5b**). We found that *NOTCH3* had higher expression in clones C and D relative to A and B as expected based on the clone-specific amplification in C&D (**Fig. 5b**). Furthermore, clones C and D had a higher VEGF pathway score, lower hypoxia score and higher HIF1A expression, a transcriptional regulator of hypoxia response (**Fig. 5b**). Interestingly, despite receiving anti-angiogenic maintenance therapy with bevacizumab, this patient still experienced disease recurrence within approximately a year of chemotherapy completion. We speculate that this genomically encoded pre-existing phenotypic state may have played a role in disease relapse due to the enhanced angiogenic potential of clone D.

In patient 009, all clones were represented in the scRNAseq and clone F was the only clone present at the final timepoint post relapse (**Fig. 5c**). We found that this clone had lower expression of JAK/STAT pathway genes, an increase in epithelial-to-mesenchymal transition (EMT) related genes including the canonical EMT marker VIM, and a lower fraction of cycling cells (**Fig. 5c**). This suggests that this clone may have an immunosuppressive phenotype, while slower cycling of cells may have rendered it less sensitive to chemotherapy. Furthermore, EMT has been associated with chemotherapy resistance^38^ and phenotypic plasticity that is permissive to developing drug resistant states^39^.

### Modeling evolutionary fitness and selection in patients

Lastly, we modeled the evolutionary properties of clonal trajectories in the context of treatment and disease recurrence. In 6/10 patients (002, 006, 014, 045, 081, 107; see **Fig. 4** & **Supplementary Figure 6**), multiple clones were detected in post-recurrence plasma samples, highlighting that chemo-resistance may be polyclonal in many patients. In 4 of these cases (009, 045, 081 and 107), clones that were not detected in cfDNA or were detected at minor frequencies at baseline became dominant at first recurrence, suggesting that although multiple clones may become chemo-resistant, some have relative fitness advantages in the context of treatment (**Fig. 4** & **Supplementary Figure 6**). Notably, while the presence of multiple resistant clones was a common observation, the overall diversity, as quantified by Shannon entropy, decreased in the final timepoint relative to baseline in 8/9 cases (p=0.027, t-test, **Supplementary Figure 6h**). This potentially reflects clone eradication during front-line treatment (surgery and chemotherapy), and that only a fraction of clones present at diagnosis comprised relapsed disease. The number of clones detected also decreased at the final time point relative to baseline (p=0.086, t-test, **Supplementary Figure 6i**).

We then tested whether changes in clonal composition could be explained by a neutral evolutionary model or whether differential fitness between clones was a more plausible explanation. We developed a Wright-Fisher^40^ population genetics based simulation and hypothesis testing framework that incorporates patient specific measurements. The simulation includes a varying population size empirically informed by CA-125 levels to model population bottlenecks due to treatment, and uses the inferred clone frequencies at baseline as starting conditions (**Fig. 6a**). Clonal trajectories were then simulated assuming neutrality (no fitness difference between clones), and the distribution of frequencies over 1000 simulations were compared to observed frequencies at the final time point to derive a p-value encoding whether the observed data is consistent with the neutral model (**Fig. 6a**). Examples of patient clone trajectories inconsistent with a neutral model include 045 and 009 (p<0.05 for at least one clone), while data from 014 could be explained with a neutral model (**Fig. 6b**). Overall, 7/10 patients had at least 1 clone whose change in frequency at the final timepoint compared to baseline could not be explained by a neutral model (**Fig. 6c**), suggesting that positive clonal selection induced by treatment may indeed be a common feature in HGSOC.

**Figure 6.**
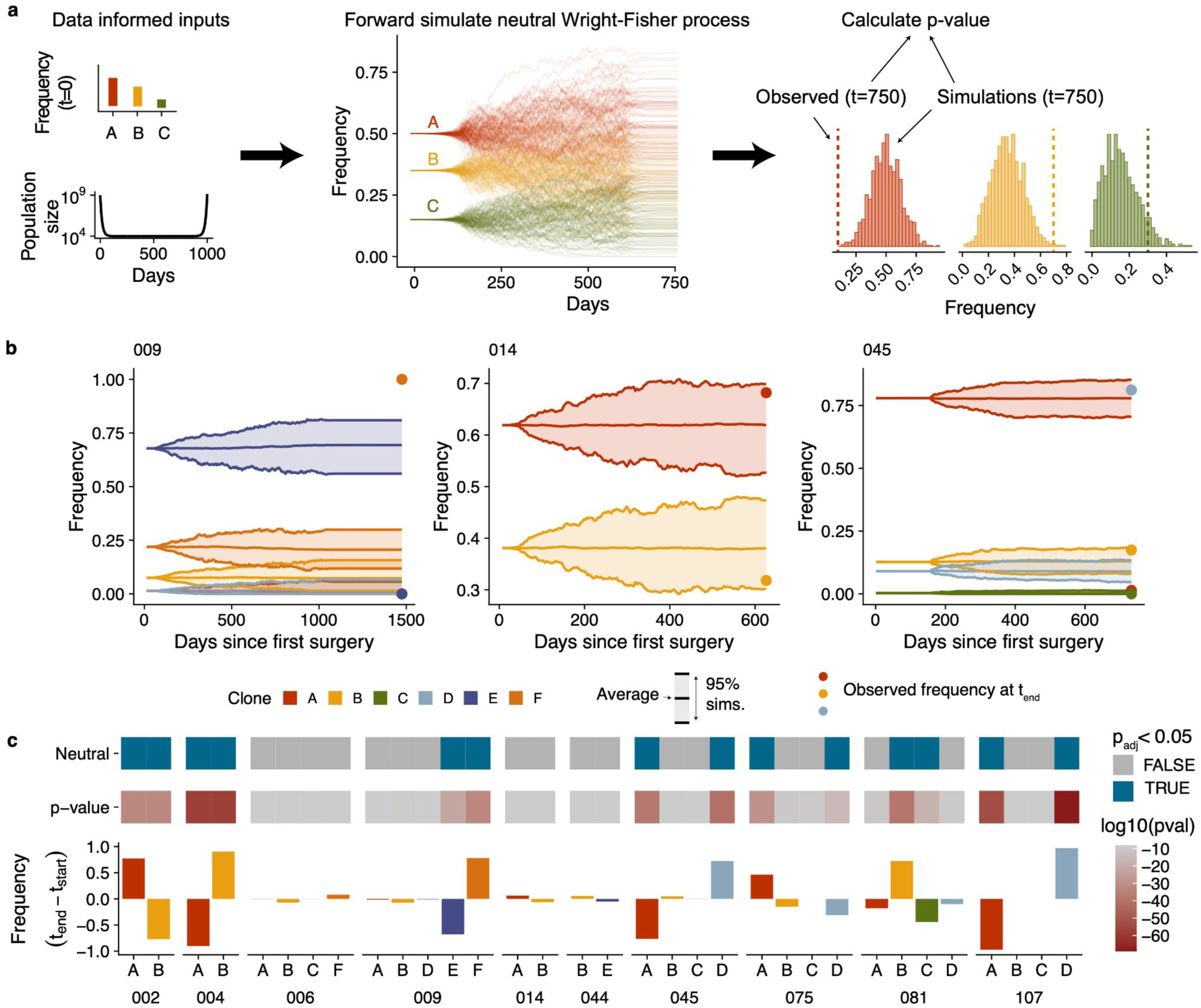
Wright-Fisher modeling. **a)** Summary of approach used to accept/reject neutrality. Frequency of clones at baseline and changes in cancer cell population informed by CA-125 levels are used as input to a neutral wright-fisher model with varying population sizes. For each sample, 1000 simulations are generated and then the distribution of frequencies at the final time point are compared to observed values. **b)** Example simulated trajectories and observed frequencies for 3 patients: 009, 014 and 045. 009 and 045 have clones that deviate from the expectations in a neutral model, while clones in 014 are consistent with a neutral model. **c)** Summary of the results of the Wright-Fisher simulation based test in 10 patients. From bottom to top: change in clone frequencies between baseline and the final timepoint which had evidence of ctDNA (in most cases the final timepoint samples), p-values per clone, neutral/non-neutral classification based on a cutoff of p(adjusted) < 0.05.

Taken together we show that clone-specific SVs can be used to track clone trajectories over time in cfDNA and that while drug resistance is often polyclonal, changes in clone frequencies are likely the result of differential fitness between clones in the context of treatment.

## DISCUSSION

Here we show that tracking clonal evolution of drug resistance is tractable in cancer patients. This was facilitated by probing structural variants in timeseries cfDNA as highly specific genomic features to monitor and model clonal evolution. Applying this approach to 10 recurrent HGSOC patients with longitudinally collected cfDNA samples we found that in many cases, clonal composition changed between diagnosis and recurrence. In most cases, drug resistance was polyclonal, but generally contained a dominant clone with frequency > 50%. Interestingly, dominant clones were typically rare at diagnosis, suggesting therapy induced selection and supported by Wright-Fisher modeling. It is noteworthy that Wright-Fisher modeling, which predicted reproducible positive selection in replicate PDX models^41^ here shows consistent properties in patients, providing motivating examples for development of predictive models. We recognize that our simulation framework neglects any spatial component whereby some clones may reside in a privileged site with different immunological properties or drug localization propensities.

Our conclusions differ from Smith *et al*^42^ which found limited copy number differences between diagnosis and recurrence samples in HGSOC. We speculate that the higher resolution single-cell measurements employed in this study are more suitable for characterizing genomic differences in such a heterogeneous disease, and that ctDNA provides a more unbiased view of the disease state compared to single-site bulk measurements. However, larger scale studies across spatio-temporal measurements may be needed to fully establish these properties.

While our study is underpowered to identify recurrent genetic features of drug resistant clones, we do observe some plausibly important features that will require additional study. These include clone-specific high-level amplifications in drug resistant clones such as *RAB25*, *ERBB2*, *CCNE1* and *NOTCH3*. Such oncogenes have targeted therapies available clinically or in development, raising the possibility that treatment could be modified adaptively based on longitudinal measurements of clone fractions and their genomic features. We also observed notable phenotypes in drug resistant clones in a subset of patients with matched scRNAseq such as upregulation of EMT and VEGF, downregulation of JAK-STAT and lower proliferation. This suggests pre-existing phenotypic states may play a role in differential treatment-sensitivity between clones. Our study confirms that drug resistance is heterogeneous and highly patient specific. For example, *CCNE1* amplification, an established indicator of poor prognosis^34^ was not a deterministic predictor of clone fitness in one of the patients. This argues for developing personalized adaptive approaches to control drug resistant clones^43–45^.

We note that the granularity of temporal sampling in this study limits the ability to accurately time emergence of drug-resistant clones. Some clone trajectories coincided with therapy modulation. We recognize this is confounded by the timing of sampling at relapse and consequently makes it challenging to establish a causal link between selective sweeps of clones and change of therapeutic selective pressure. In future studies, more granular cfDNA sampling at regular intervals would address this interpretation challenge, but our data is nevertheless indicative of clone-specific selective sweeps on switch of therapy.

Our study is motivated by exploiting structural variants as specific endogenous clonal markers for evolutionary tracking. While we focused on HGSOC, we expect our approach will generalise to any tumor type with i) polyclonal disease and ii) with characteristic genomic instability such as triple negative breast, osteosarcoma, high grade endometrial, esophageal, diffuse gastric and *EGFR*-mutant non-small cell lung cancer^21^.

Finally, we expect that drug resistance in HGSOC and other cancers to be multi-factorial, with both transcriptional and epigenetic plasticity operating in tandem with pre-adapted and genomically encoded phenotypic states. We contend that the framework we establish here is poised to quantify the proportion of drug resistance that is explained by therapeutic selective pressure and can inform future evolution-informed adaptive clinical trials.

## Methods

### Sample collection

All enrolled patients were consented to an institutional biospecimen banking protocol and MSK-IMPACT testing^46^, and all analyses were performed per a biospecimen research protocol. All protocols were approved by the Institutional Review Board (IRB) of Memorial Sloan Kettering Cancer Center. Patients were consented following the IRB-approved standard operating procedures for informed consent. Written informed consent was obtained from all patients before conducting any study-related procedures. The study was conducted in accordance with the Declaration of Helsinki and the Good Clinical Practice guidelines (GCP). We collected fresh tumor tissues at the time of upfront diagnostic laparoscopy or debulking surgery, as described in McPherson et al.^22^. Blood collection was carried out longitudinally over a five-year period (2019-2024). Two Streck tubes for cfDNA were collected in each visit. If possible, blood was collected in the Outpatient Clinic at Memorial Sloan Kettering Cancer Center. Alternatively, blood samples were collected in the operating room when patients were undergoing debulking surgery or laparoscopy.

### Sample processing

Streck tubes were submitted to the MSK laboratory medicine facility after collection and processed for plasma and buffy coat separation, as well as DNA extraction.

### Clinical data

In this cohort study, we extracted clinical annotations from electronic health records of 19 patients treated at Memorial Sloan Kettering Cancer Center for HGSOC. For these patients we collected contemporaneous longitudinal data from their initial HGSOC diagnosis as well as historical data, if available. Clinical data included laboratory measurements, surgical procedures and medications. CA-125 measurements were obtained as part of patients’ routine clinical care from blood samples collected at baseline, during therapy and subsequent follow up visits. All dates are relative to the time of first surgery for each patient, ie day 0 is the date of primary debulking or laparoscopic biopsy.

### Recurrence data

Recurrence dates are defined by “progression of disease” (POD), a patient without improvement after treatment or while on maintenance therapy based on CT scan. Improvement or lack thereof is determined based on CT scan impressions (e.g. an increase in a lymph node or unchanged tumor implants). We define patients as “alive with disease” (AWD) if they have not achieved remission but have also opted out of new treatment lines and/or are on observation.

### DLP single cell whole genome sequencing processing

The mondrian single cell whole genome sequencing suite of tools and pipeline was used for processing of the single cell whole genome sequencing. This single cell whole genome sequencing dataset is a subset of the dataset used in McPherson *et al.*^22^, see this publication for full details of the data generation and processing. We describe in brief the processing here. Sequencing reads were aligned to hg19 using BWA-MEM. Read counts were calculated in 500kb bins across the genome and GC-corrected, these values were input into HMMcopy to infer integer copy number (ranging from 0-11). We then applied the cell quality classifier described in Laks et al and removed any cells with quality < 0.75. In addition we removed replicating cells, multiplet cells and cells suspected to be the result of multipolar divisions, see McPherson *et al.*^22^ for a detailed description of the filtering criteria. We then applied SIGNALS v0.7.6 to infer haplotype specific copy number using default parameters.

### SNV calling in DLP

To detect SNVs in each dataset, reads from all cells from a DLP+ patient were merged to form ‘pseudobulk’ bam files. SNV calling was performed on these libraries individually using mutect. A panel of normals was constructed by identifying normal cells from every patient, merging them and then running the mutect2 panel of normal option. Mutect2 filter was used to filter variants. We then ran Articull (manuscript in prep.) to remove artifacts that are specific to DLP+ due to the shorter than average insert size. This filtered set of variant were then genotyped in individual cells using cellSNP v.1.2.2^47^.

### Structural variant calling in DLP

To detect SVs in each dataset, we also used the merged pseudobulk bam files. LUMPY^48^ and deStruct^49^ were run on these pseudobulk libraries. Events were retained if they were detected in deStruct and could be matched in the LUMPY calls. Breakpoint predictions were considered matched if the positions involved were each no more than 200 nucleotides apart on the genome and the orientation was consistent.

SVs called in the pseudobulk library were then genotyped in single cells. To do this we used a modified version of SVtyper^50^ (available at https://github.com/marcjwilliams1/svtyper). One key modification was rounding the read count up rather down, the read count computation internally in SVtyper are MAPQ scores rather than counts so are non-integers before rounding and outputting to a vcf. This change is necessary in single cells as typically we observe only a single read supporting a SV. SVtyper computes the number of reads that support the reference (these are reads that directly span the genome reference at the breakpoint locations) and the number of reads that support the alternate allele. Alternate allele counts are either split reads that directly sequence the breakpoint or discordant reads that have larger than expected insert sizes or align to different chromosomes in the case of translocations. Clipped reads that support the breakpoint are also computed, to be more conservative we did not include these reads in the total of SV supporting reads. We made an additional modification requiring split reads to match both sides of the breakpoint to contribute to read counts, in the default version, a split read aligning to one side of the breakpoint would contribute 0.5 counts. This option is available via the `–both-sides` command line option.

### Phylogenetic inference and clone assignments

MEDICC2 was used to infer phylogenetic trees using haplotype specific copy number as input, see McPherson *et al*^22^ for further details. We then manually identified clades in the tree that were the ancestor of clade specific genomic features of interest. These included whole genome doubling, whole chromosome and chromosome arm gains or losses and focal amplifications. Clones were then defined as the set of cells that were descendents of each clade of interest. Clones with their genomic features of interest can be found in supplementary table 5.

### Clone level 10kb resolution copy number calling

Once cells were assigned to clones we additionally called integer copy number at 10kb resolution at the clone level. Read counts were computed in 10kb bins across the genome in every cell and then summed across cells assigned to each clone. Aggregated read counts were then normalized against the read counts from any normal diploid cells sequenced in the same library and then GC corrected using the same modal GC correction described in Laks *et al*^17^. These normalized GC corrected read counts were then adjusted for ploidy of the clone and then we applied a Hidden Markov Model to compute integer read counts. The HMM model (code available at https://github.com/shahcompbio/HMMclone) uses a state space of 0-15 with each state assumed to be a normal distribution with standard deviation 0.2 and mean equal to the integer copy number. The standard deviation was determined empirically from the data. The viterbi algorithm was used to compute the most likely copy number profile.

### Assigning SVs to clades/clones

Assigning SVs to clones was done using the matrix of read counts per SV per cell, the cell to clone label mapping, and the clone level 10kb copy number profiles. First, we summed the SV supporting reads across clones giving an SV by clone matrix. Any SVs with non-zero read counts were assumed to be present in the clone. In addition, when SVs could be mapped to copy number changepoints identified at 10kb resolution, we additionally checked whether there existed other clones that had the same copy number changepoint but lacked read level support for the SV. In these cases, the SV was also assumed to present in that clone. This was to circumvent cases where the total number of cells was too low to confidently assume the absence of a particular SV. In some cases, no SVs could be found that were specific to a clone, this was largely due to clones being too small and consequently lacking the cumulative sequencing coverage to detect SVs in pseudobulks. In such cases we used coarser clone definitions comprising a larger number of cells.

### Bulk whole genome sequencing and MSK-IMPACT

The bulk whole genome sequencing and MSK-IMPACT targeted sequencing was originally published in Vázquez-García *et al.* ^23^. See this publication for data generation, processing and data access.

### Probe design and synthesis

For most patients we identified 1,000 genomic features encompassing structural variants, single nucleotide variants and germline SNPs. For samples that constituted our pilot (patients 068, 065, 044, 003 and 026) the number of features was lower, between 250 and 400 and included only a limited number of SNVs. Within the SV and SNV groups these could be classed into Clonal (present in every tumor cell) or Subclonal (present in a fraction of tumor cells). The number of probes from each class was variable between patients due to differences in the number of SVs and SNVs called in each patient as well as the clonal structure in each patient. We first required 200 clonal SVs and 200 clonal SNVs. The remaining 600 probes were split between subclonal SVs and SNVs. We ensured we had 200 subclonal SNVs and then the remaining slots were given to subclonal SVs, if there were still slots remaining then we included additional subclonal SNVs. Within the SNV class we included any SNV annotated as “High Impact” in the MSK-IMPACT targeted sequencing. Probes were synthesized by IDT (Integrated DNA Technologies) using the xGen MRD hybrid probes, from 120bp sequences provided as FASTA files. A small panel of germline SNPs were also included in order to provide a means to identify sample swaps that may inadvertently occur during sample preparation but were not needed.

### cfDNA duplex sequencing analysis

We used the MSK-ACCESS protocol to generate the sequencing data, this protocol is described in detail in Rose-Brannon *et al.*^25^. The gene panel used in Rose-Brannon *et al.* was swapped for the patient specific probe sets. Patient probes from at least 2 patients were pooled together so that for each patient probe set we could estimate background error rates by looking at the counts supporting SVs and SNVs in off target patients.

To process the cfDNA sequencing we used a suite of tools developed by the Centre for Molecular Oncology informatics team at MSK for use with the MSK-ACCESS assay (https://github.com/msk-access). The nucleo pipeline was used to generate bam files from fastq files. The output of this pipeline is double strand error corrected bam files(duplex), single strand (simplex) and uncorrected bam files which can then be used for downstream applications. Read counts of supporting and reference reads for SNVs and Indels were extracted using https://github.com/msk-access/GetBaseCountsMultiSample. This takes a MAF file as input and outputs a MAF file with additional columns for the read counts in duplex, simplex or uncorrected bam files. To extract read counts for SVs we used the same version of SVtyper modified for use with DLP+ described above. We required that alignments had evidence of both sides of the breakpoint to be included (implemented in an additional SVtyper option `–both_sides`).

### Computing error rates in cfDNA

To compute error rates across sequencing types (duplex, simplex, raw uncorrected) and mutation types (structural variants and single nucleotide variants) we applied the patient specific probe set to at least one other off-target patient. We then summed the counts of reference supporting reads and variant supporting reads for off-target variants and defined the error rate as variants supporting reads divided by total number of reads. We then defined the limit of detection (LOD) per sequencing type and mutation type as twice the largest error rate seen in each class. Given we observed no errors for SVs in simplex and duplex sequences we defined the LOD as the inverse of the total number of reference supporting reads giving us an upper bound. This gives the following LOD: 8.5×10^-8^ (duplex, SV), 3.2×10^-5^ (duplex, SNV), 1.5×10^-7^ (simplex, SV), 20×10^-5^ (simplex, SNV), 8.6×10^-7^ (uncorrected, SV), 158×10^-5^ (uncorrected, SNV). LOD for combined duplex and simplex read counts is 5.4×10^-8^ for SVs and 7.3×10^-5^ for SNVs. cfDNA samples were positive for ctDNA if the total number of variant reads divided by the total number of reference reads summed across the collection of patient specific variants was greater than the LOD. Given the low error rates for both simplex and duplex SVs we used the combined read counts from both for the results reported in the main text.

### Estimating clone frequencies

To estimate clone frequencies we calculated VAFs for each clone by summing the total number of variant supporting reads and dividing them by the total number of reads for all variants assigned to a clone. We did not correct for copy number as biases in the sequencing data are likely greater than biases due to copy number (probes are constructed based on the variant sequence not wild type). Furthermore, VAFs can vary over multiple orders of magnitude due to the high sequence depth, much larger than the influence of any copy number correction. We saw highly concordant clone frequency estimates using either structural variants or single nucleotide variants, supporting this approach. To plot the changes in frequency over time we normalized VAFs so that they summed to 1 at each time point, then applied a spline function to smooth values between time points. When no tumor DNA was detected, we allowed all clones to have VAF = 0. Smoothing was done using the the `splinefun` function in R with method = "monoH.FC". This resulted in values that were greater than 1 or less than 0 in some cases, we therefore re-normalized the data so that frequencies were positive and summed to 1 at each time point. In addition, when there were large periods of time pre clinical-recurrence without cfDNA samples we assumed tumor DNA was 0 (for example in patient 107). We did not include clone frequency estimates when plasma tumor fractions were < 10^-4^ (estimates based on truncal SVs), reasoning that clone frequency estimates at such low tumor fractions would be unreliable and suffer from dropout issues. Given the low error rates, we used the uncollapsed raw sequencing for estimating clone frequencies using SVs. For estimating clone frequencies using SNVs we used the same approach for SVs but used duplex consensus sequences for read counting due to the higher error rates for SNVs.

### Identifying BRCA reversion mutations

For *BRCA1/2-*mutant cases, we also included probes that captured exonic regions within 200bp of the mutation, enabling detection of proximal *BRCA1/2* reversion mutations^51,52^. We used revmut (https://github.com/inodb/revmut) to identify putative BRCA reversion mutations in the first instance. In addition, we inspected alignments in IGV around the BRCA mutations to look for any additional putative reversion mutations not identified by revmut. This is how we found the reversion mutation present in patient 009. This mutation was a large 1.37kb deletion that excised the germline mutation, alignments with the same breakpoint sequence, aligning to the same locations were found in 3 post-recurrence samples. This mutation was likely not identified using revmut due to it being unusually large compared to previously reported BRCA reversion mutations.

### Wright-Fisher modelling and hypothesis testing

In order to test for non-neutrality in clone frequencies over time we implemented a modelling and hypothesis testing framework based on a multi-species Wright-Fisher model with varying population size. Population size was assumed to be 10^9^ at the time of surgery (t=0) and then varied according to CA-125 levels. We set the population size at the time point with the lowest CA-125 level *N*_*low*_ = 10^4^, assuming this was the period with the smallest tumor cell population. We then set the population size (N) to vary exponentially according to the following equation:

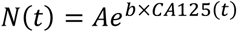

Where *A* = *N*(0) × *e*^−*b*×*CA*125(0)^ and 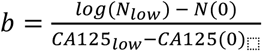. We then used the multinomial distribution to simulate clone frequencies over time:

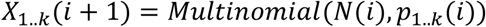

Where *X*_1..*k*_(*i* + 1) is the population size of each clone k in generation i+1, N(i) is the total population size in generation i and *p*_1..*k*_(*i*) are the clone frequencies in generation i for the k clones. For generation *i* = 1, *N*(*i* = 1) = 10^9^ and *p*_1..*k*_(*i* = 1) are given by the clone frequencies estimated from cfDNA at t=0. We then forward simulate this process for the clinical timecourse of each individual patient 1000 times giving a distribution of clone frequencies at *t*_*end*_. We assumed a generation time of 4 days and *t*_*end*_ was set to be the final cfDNA timepoint in each patient. We then calculated a z-score, comparing the observed clone frequency from data to the mean and standard deviation of the simulated frequencies in order to calculate a p-value for each clone under the hypothesis of neutral evolution.

### cfDNA whole genome sequencing

Whole genome libraries constructed during the duplex sequencing assay library prep were sequenced to 20X on an illumina NovaSeq using 100bp reads. Reads were mapped to hg19 using BWA-MEM^53^. Read counts in 100kb bins across the genome were calculated and GC corrected using QDNAseq^54^. In order to compare this to data from the duplex sequencing targeted assay we used information from the DLP copy number profiles and the clone fractions inferred from the hybrid capture targeted sequencing assay to predict what these copy number profiles should look like. Copy number ratio (*R*) in bin *i* are given by:

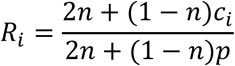

Where *n* is the normal fraction, *c*_*i*_ is the copy number in bin *i* and *p* is the ploidy of the tumor. We know *n* from the TP53 VAF in cfDNA, for *c*_*i*_ and *p* we took the weighted average across clones, with weights given by the estimated clone fractions at each time point.

### scRNAseq data generation and processing

The scRNAseq data was originally published in Vazquez-Garcia et al.^23^, full details of the processing can be found here. Pathway scoring was performed with PROGENY^55^ or the Seurat module scoring function using hallmark pathways.

### TreeAlign

To match scRNAseq cells to clones identified in DLP we used TreeAlign^37^. To do this, we genotyped the same set of heterozygous SNPs used to call allele specific copy number in DLP+ in scRNAseq using cellSNP^47^. The per cell SNP count matrix was then input into TreeAlign along with clone assignments and 10kb clone copy number profiles derived from DLP. We used the CloneAlignClone method and used default parameter values apart from min_clone_assign_prob = 0.5. scRNAseq data was available for patients 107, 014, 045, 009 and 002, however following application of TreeAlign in some patients, clones were represented minimally due to differences in data collection from different sites. In patients 107 and 009, all clones were represented with at least 100 cells present from each clone, we therefore focussed on these cases when comparing drug resistant to drug sensitive clones. To compare transcriptional heterogeneity for each clone we took the mean value of the per cell seurat derived module scores or progeny scores per clone, then for each patient calculated the maximum value minus the minimum value. These per patient max-min values were then plotted as violin plots ordered by the average difference across the cohort of patients.

### Data organization

To facilitate integration of data across multiple modalities we used the isabl platform^56^. Isabl is a databasing and data access platform which allows users to straightforwardly link multiple datasets from the same patient and chain together pipelines across modalities.

## Data availability

Summary tables include sequencing coverage, cfDNA tumor fractions, clone frequencies from SVs and SNVs, genomic features of defined clones and error rates per patient. Raw sequencing data will be available in dbGAP upon publication. Processed copy number calls and variant read counts in cfDNA will be available in Synapse (accession number syn25569736).

## Code availability

The pipeline to process DLP+ scWGS is available at https://github.com/mondrian-scwgs. SIGNALS^18^ was used for most plotting and scWGS analysis and is available at https://github.com/shahcompbio/signals. Clone copy number profiles at 10kb were computed using HMMClone (https://github.com/shahcompbio/HMMclone). The modified version of SVtyper for use with single cells and hybrid capture duplex sequencing is available at https://github.com/marcjwilliams1/svtyper.

## Acknowledgements

This project was funded in part by Cycle for Survival supporting Memorial Sloan Kettering Cancer Center and the Halvorsen Center for Computational Oncology. S.P.S. holds the Nicholls Biondi Chair in Computational Oncology and is a Susan G. Komen Scholar. This work was funded in part by Break Through Cancer and by awards from the Ovarian Cancer Research Alliance (OCRA) Collaborative Research Development Grant [648007] and NIH R01 CA281928-01 to S.P.S., OCRA Ann Schreiber Mentored Investigator Award to I.V.-G. [650687], NCI pathway to independence award M.J.W [K99CA256508], OCRA Liz Tilberis Award to D.Z., the Department of Defense Congressionally Directed Medical Research Programs to S.P.S., D.Z. and B.W [W81XWH-20-1-0565], the Seidenberg Family Foundation, the Cancer Research UK Cancer Grand Challenges Program to S.P.S. [C42358/A27460], NIH U24 [CA264028] to S.P.S., the Marie-Josée and Henry R. Kravis Center for Molecular Oncology and the National Cancer Institute (NCI) Cancer Center Core Grant [P30-CA008748]. B.W. is funded in part by Breast Cancer Research Foundation and NIH/NCI P50 CA247749 01 grants. D.Z. is funded by NIH grant R01 CA269382.

## Competing interests

B.W. reports grant funding by Repare Therapeutics paid to the institution, outside the submitted work, and employment of a direct family member at AstraZeneca. C.A. reports grants from Clovis, Genentech, AbbVie and AstraZeneca and personal fees from Tesaro, Eisai/Merck, Mersana Therapeutics, Roche/Genentech, Abbvie, AstraZeneca/Merck and Repare Therapeutics, outside the scope of the submitted work. C.A. reports clinical trial funding to the institution from Abbvie, AstraZeneca, and Genentech/Roche; participation on a data safety monitoring board or advisory board in AstraZeneca and Merck; unpaid membership of the GOG Foundation Board of Directors and the NRG Oncology Board of Directors. M.F.B reports consulting fees (Eli Lilly, AstraZeneca, Paige.AI), Research Support (Boundless Bio) and Intellectual Property Rights (SOPHiA Genetics). BL reports Intellectual propery rights (SOPHiA Genetics) and licensing royalties (BioLegend/Revvity). C.F. reports research funding to the institution from Merck, AstraZeneca, Genentech/Roche, Bristol Myers Squibb, and Daiichi; uncompensated membership of a scientific advisory board for Merck and Genentech; and is a consultant for OncLive, Aptitude Health, Bristol Myers Squibb and Seagen, all outside the scope of this manuscript. D.S.C. reports membership of the medical advisory board of Verthermia Acquio Inc and Biom’up, is a paid speaker for AstraZeneca, and holds stock of Doximity, Moderna, and BioNTech. D.Z. reports institutional grants from Merck, Genentech, AstraZeneca, Plexxikon, and Synthekine, and personal fees from AstraZeneca, Xencor, Memgen, Takeda, Astellas, Immunos, Tessa Therapeutics, Miltenyi, and Calidi Biotherapeutics. D.Z. own a patent on use of oncolytic Newcastle Disease Virus for cancer therapy. N.A.-R. reports grants to the institution from Stryker/Novadaq and GRAIL, outside the submitted work. S.P.S. reports research funding from AstraZeneca and Bristol Myers Squibb, outside the scope of this work.

## Figures

**Supplementary Figure 1 Swimmer plot**

**Supplementary Figure 2 SV + DLP summary**

**Supplementary Figure 3 Study summary**

**Supplementary Figure 4 10kb vs 500kb copy number profiles**

**Supplementary Figure 5 Copy number heatmaps**

**Supplementary Figure 6 Clonal evolution trajectories**

**Supplementary Figure 7 Validation using WGS of plasma**

## Tables

**Table S1 - cfDNA coverage statistics**

**Table S2 - cfDNA tumor fractions**

**Table S3 - Clone frequencies**

**Table S4 - Genomic features of clones**

**Table S5 - Error rates**

---

**Supplementary Figure 1.**
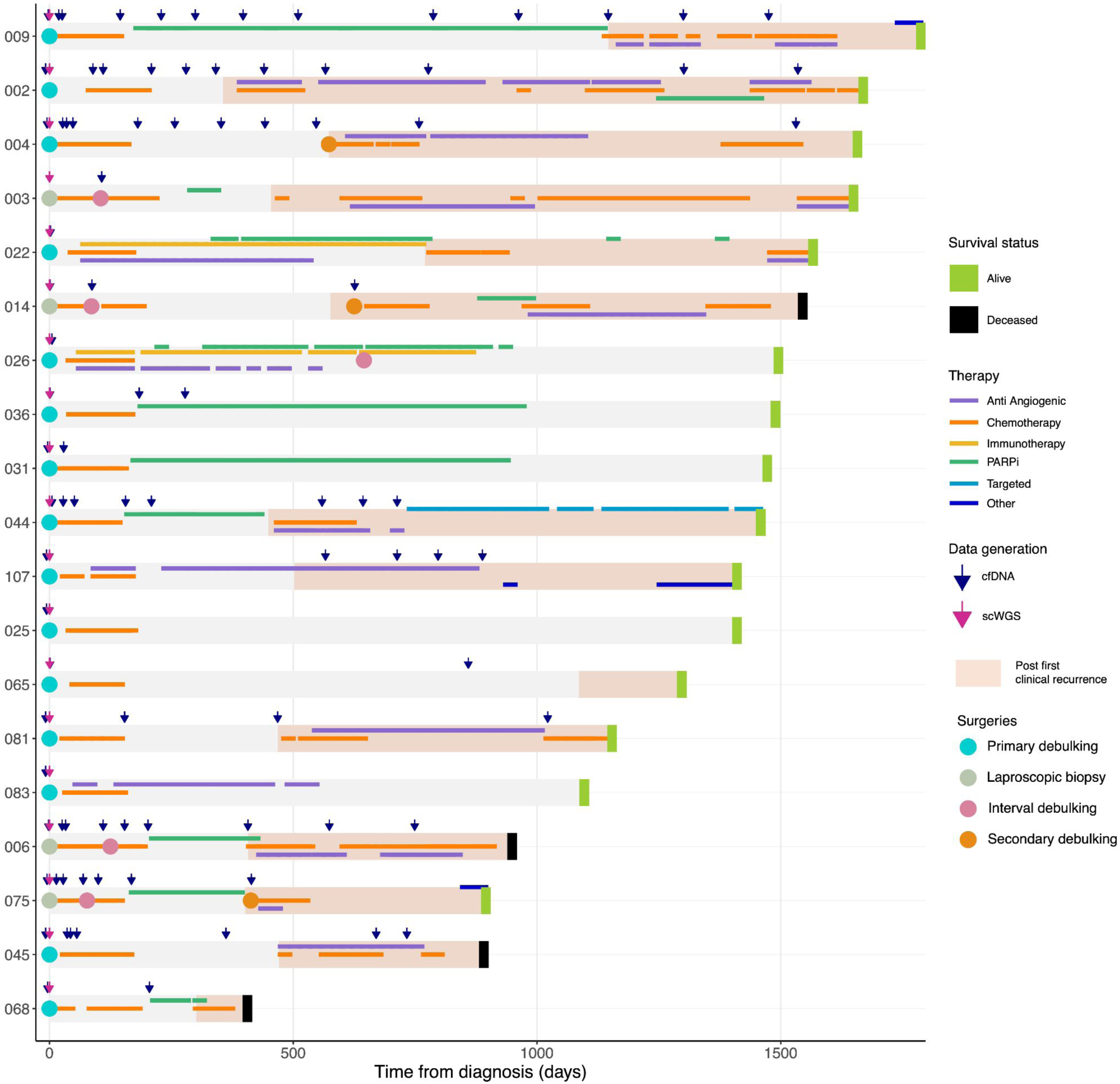
Swimmer plot showing clinical history of all 19 patients included in the study. Shown are survival status, therapies, surgeries time of first clinical recurrence and data generation timepoints. Days are relative to day of first surgery, ie Day 0 is the date of primary debulking or laparoscopic biopsy.

**Supplementary Figure 2.**
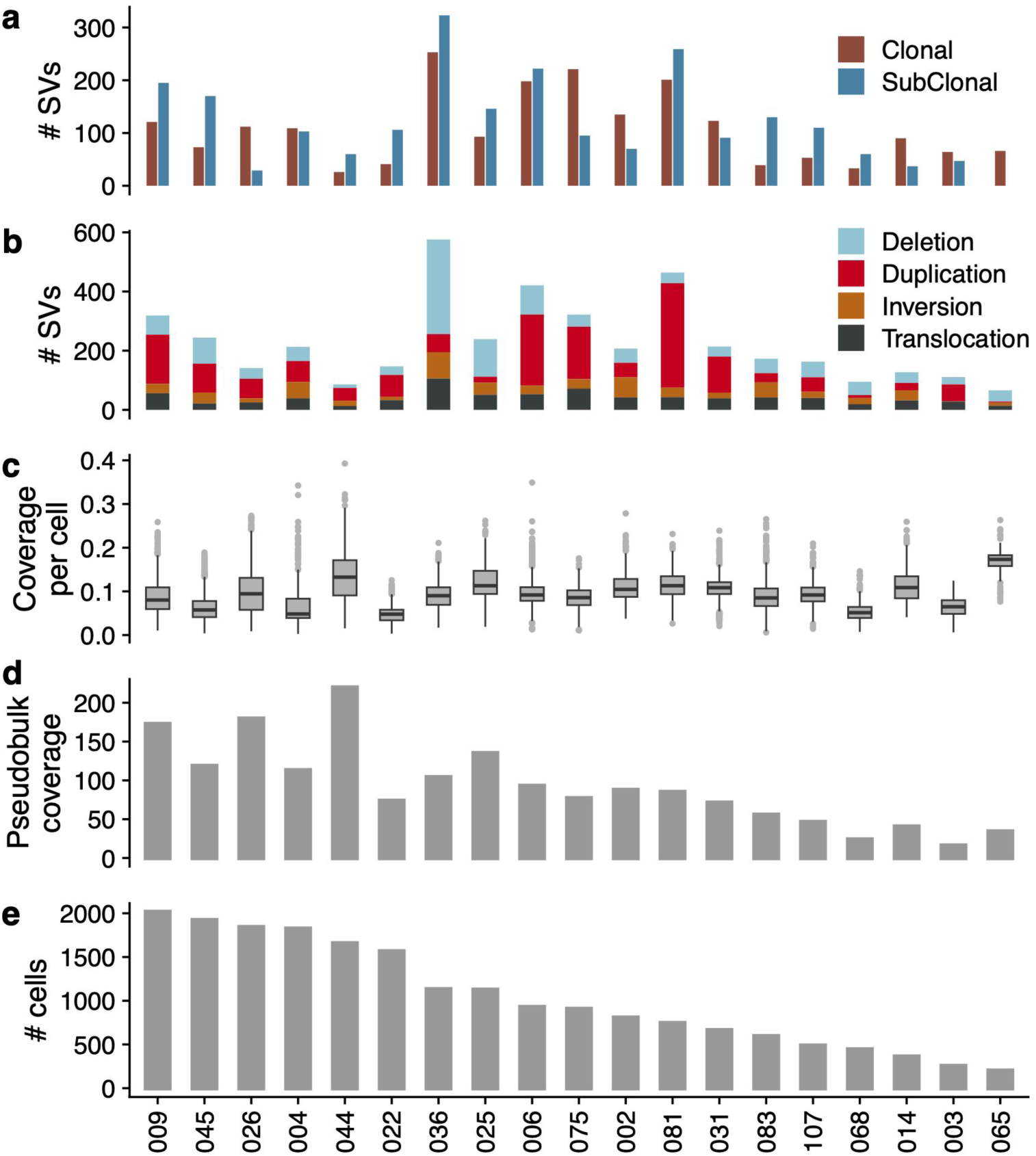
a) Number of clonal and subclonal SVs per patient b) Total number of SVs called per patient by SV type c) Distribution of coverage per cell per patient d) Pseudobulk coverage per cell (summed coverage across all cells) e) Number of high quality cells per patient

**Supplementary Figure 3.**
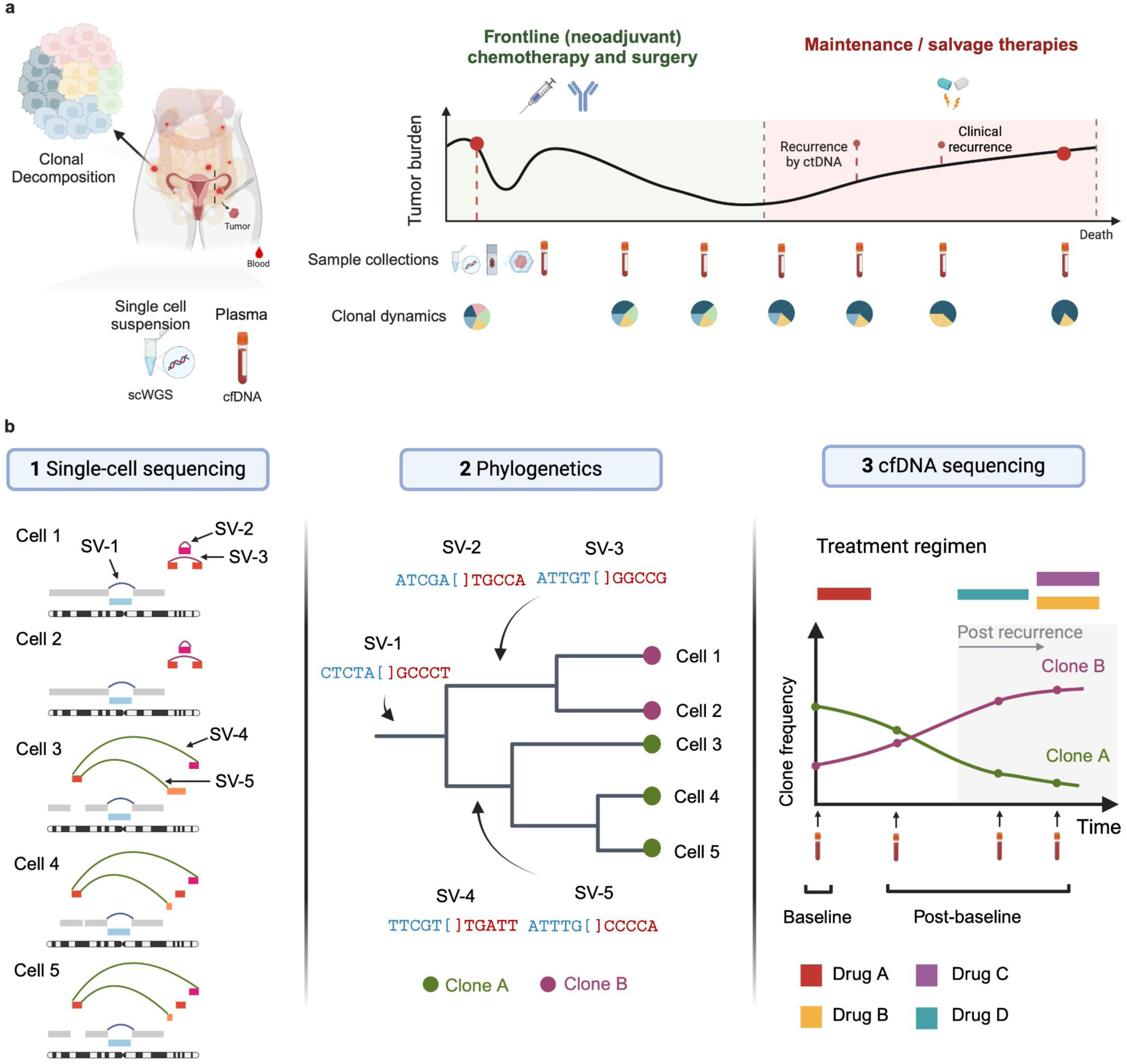
a) Study summary, showing typical clinical history of HGSOC patient, specimen sample collection protocol. b) Workflow showing clonal evolution tracking using structural variants identified in single-cell whole genome sequencing and assigned to clones using single-cell phylogenetics. These clone specific SVs are then followed in cfDNA using deep duplex error corrected sequencing.

**Supplementary Figure 4.**
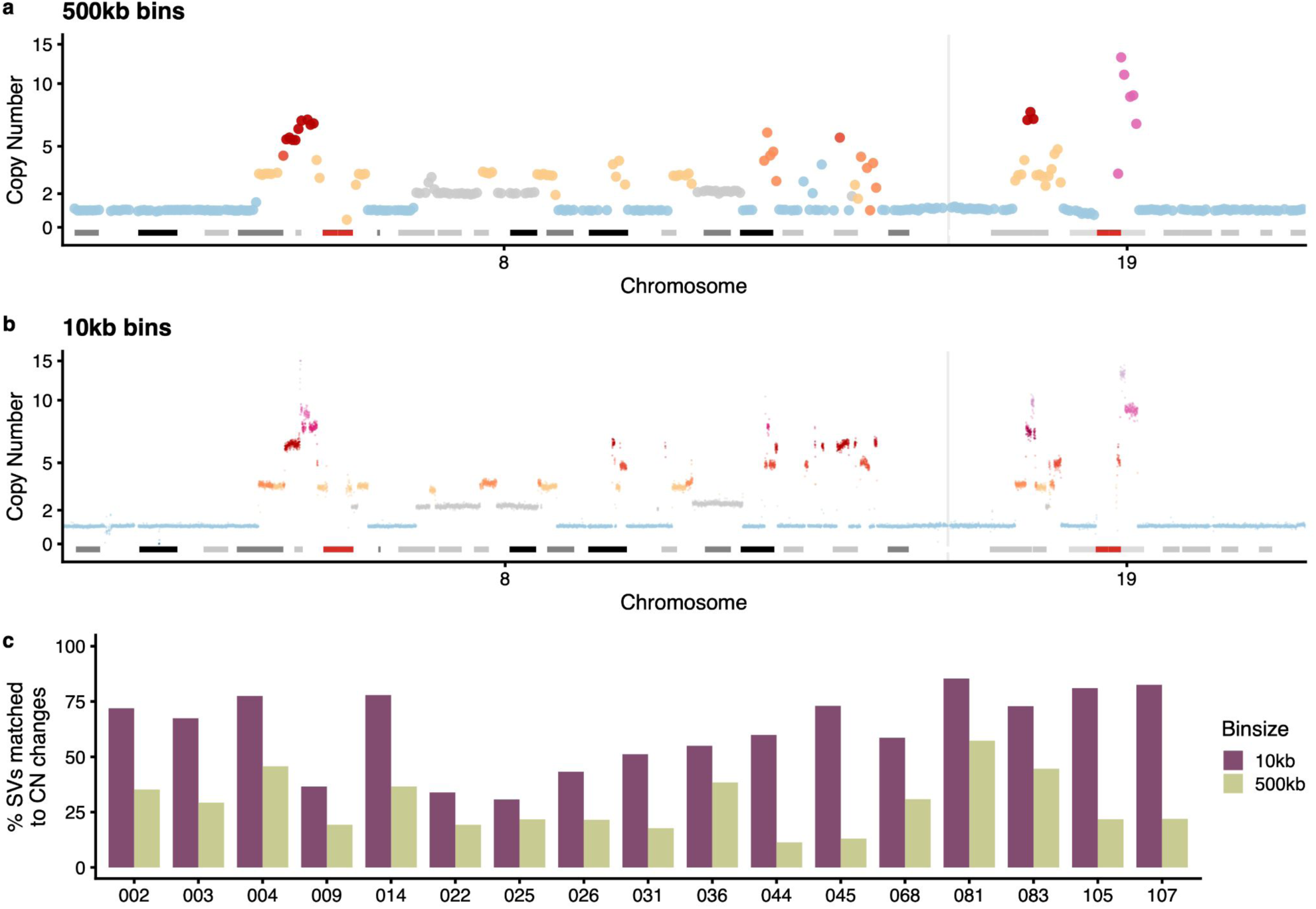
Copy number plots of chromosome 8 and 19 from OV-004 using 500kb bins a) and 10kb bins b). c) proportion of SVs that could be matched to copy number transitions at 10kb and 500kb bins

**Supplementary Figure 5.**
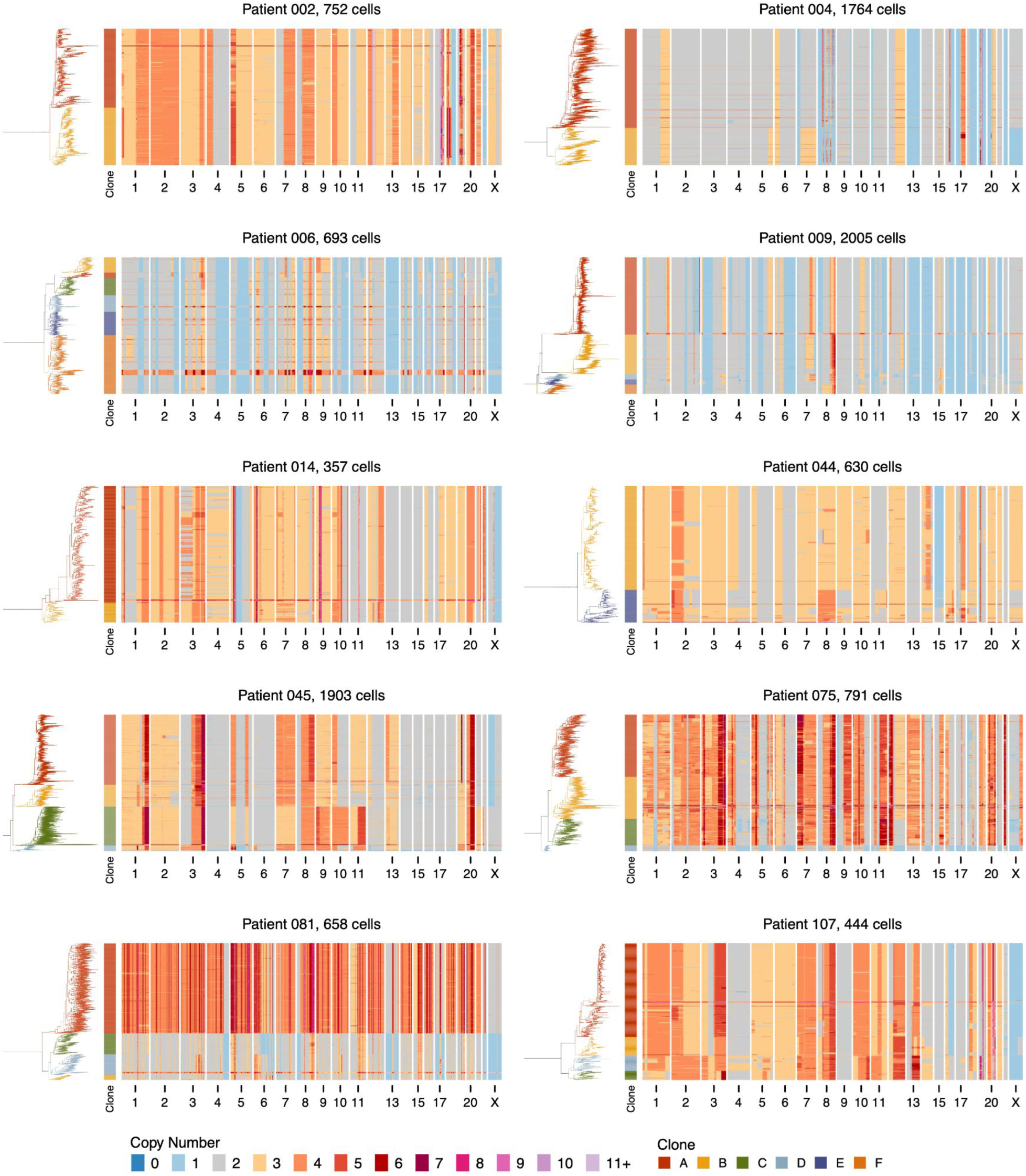
scWGS copy number heatmaps and phylogenetic trees for the 10 patients with longitudinal tracking data. The title of each plot gives the patient ID and the total number of cells. Each row shows the copy number profile of a cells, rows are ordered by the MEDICC2 derived phylogenetic tree shown on the left of each plot. Trees are coloured by clone assignments.

**Supplementary Figure 6.**
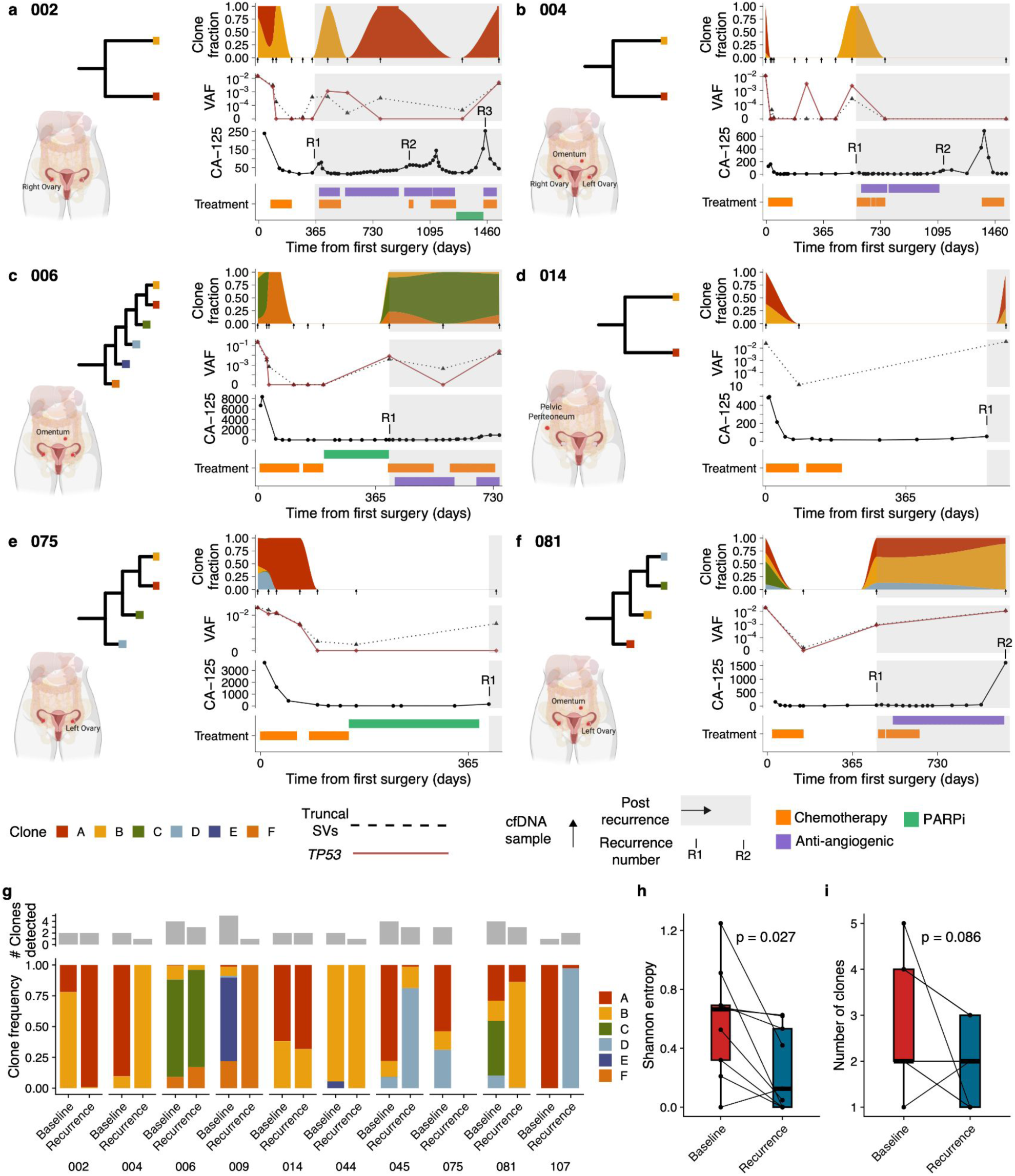
Clonal evolution tracking in 6 patients a)-f). For each patient we show the anatomical sites sequenced with DLP, a phylogenetic tree of the clones, then clonal fractions, mean truncal SV VAF and TP53 VAF, CA-125 and treatment history over time. g) Summary of the clonal composition at baseline and recurrence (final time point if more than one post-recurrence time point) for 9 patients. h) Distribution of shannon entropy at baseline and recurrence i) Number of clones detected at baseline and recurrence.

**Supplementary Figure 7.**
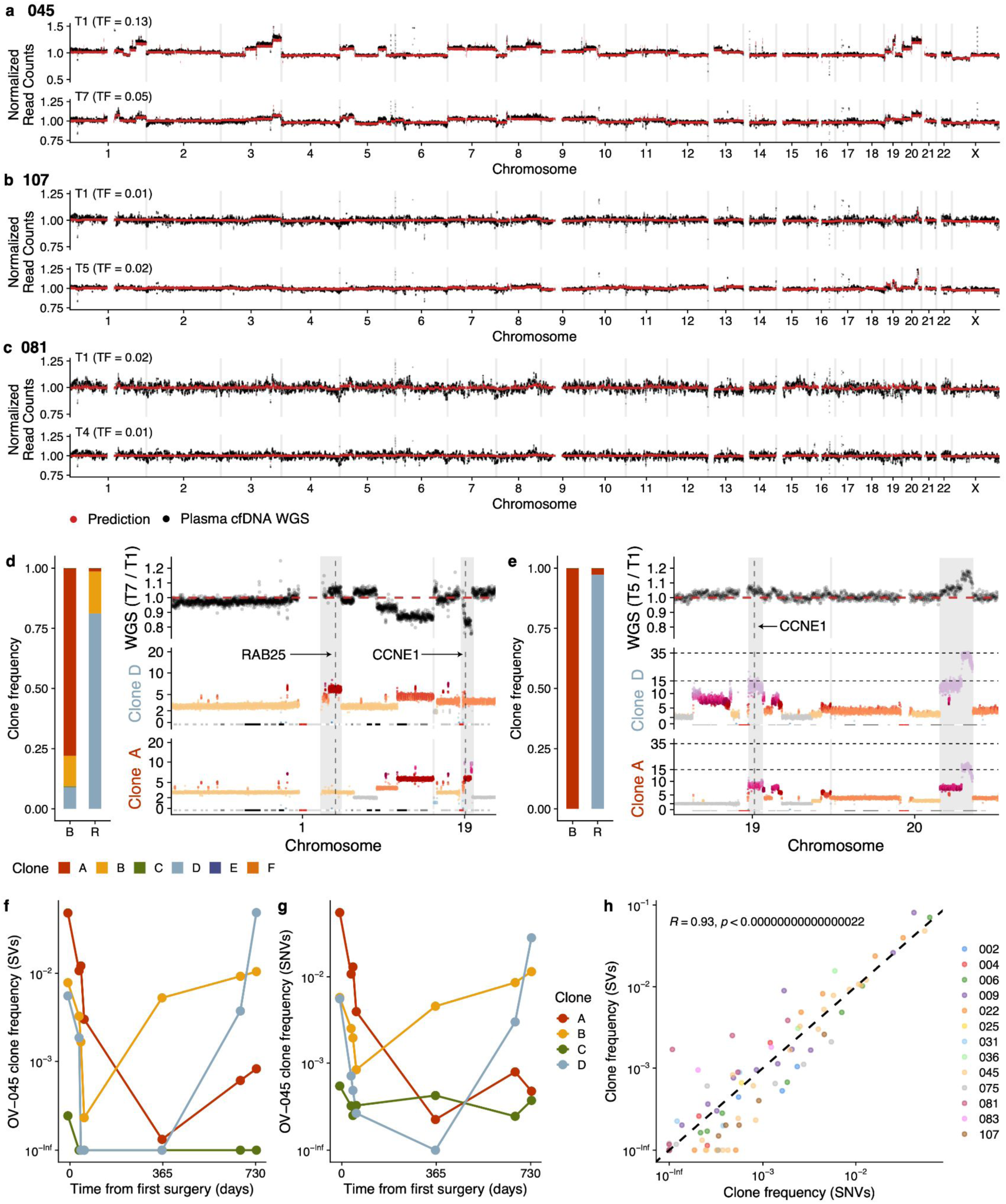
Normalized read counts at baseline and recurrence from whole-genome sequencing of cfDNA from 3 patients a)-c). Black dots are the data, red dots are predictions based on copy number profiles from DLP and inferred tumor and clone fractions from targeted sequencing. The text above each plot denotes the time point and the tumor fraction (TF) based on TP53 mutation. d) Zoom in on regions with high level amplifications in patients 045, left hand bar plots show the clone fractions at T1 and T7 then right hand side show copy number profiles of 2 most abundant clones from DLP at the bottom and ratio of normalized read counts of plasma WGS at T7 vs T1. Shaded areas highlight copy number amplification specific to one of the clones. e) Zoom in on regions with high level amplifications in patient 107. Clone frequencies over time calculated from SVs (f) and SNVs (g) for patient OV-045. c) Scatter plot of all clone frequencies calculated using SNVs and SVs, dashed line indicates y-x line. Included in this plot are clone frequency estimates from samples with purity > 0.1% and clones with at least 4 SVs and SNVs.

## References

1. Kurta, M. L. et al. Prognosis and conditional disease-free survival among patients with ovarian cancer. J. Clin. Oncol. 32, 4102–4112 (2014).

2. Siegel, R. L., Giaquinto, A. N. & Jemal, A. Cancer statistics, 2024. CA Cancer J. Clin. 74, 12– 49 (2024).

3. Black, J. R. M. & McGranahan, N. Genetic and non-genetic clonal diversity in cancer evolution. Nat. Rev. Cancer (2021) doi:10.1038/s41568-021-00336-2.

4. Ingles Garces, A. H., Porta, N., Graham, T. A. & Banerji, U. Clinical trial designs for evaluating and exploiting cancer evolution. Cancer Treat. Rev. 118, 102583 (2023).

5. Lan, X. et al. Fate mapping of human glioblastoma reveals an invariant stem cell hierarchy. Nature 549, 227–232 (2017).

6. Yang, D. et al. Lineage tracing reveals the phylodynamics, plasticity, and paths of tumor evolution. Cell (2022) doi:10.1016/j.cell.2022.04.015.

7. Bhang, H.-E. C. et al. Studying clonal dynamics in response to cancer therapy using high-complexity barcoding. Nat. Med. 21, 440–448 (2015).

8. Wan, J. C. M., et al. Liquid biopsies for residual disease and recurrence. Med (N Y) 2, 1292– 1313 (2021).

9. Cescon, D. W., Bratman, S. V., Chan, S. M. & Siu, L. L. Circulating tumor DNA and liquid biopsy in oncology. Nature Cancer 1, 276–290 (2020).

10. Ignatiadis, M., Sledge, G. W. & Jeffrey, S. S. Liquid biopsy enters the clinic - implementation issues and future challenges. Nat. Rev. Clin. Oncol. 18, 297–312 (2021).

11. Abbosh, C. et al. Tracking early lung cancer metastatic dissemination in TRACERx using ctDNA. Nature 616, 553–562 (2023).

12. Abbosh, C. et al. Phylogenetic ctDNA analysis depicts early-stage lung cancer evolution. Nature 545, 446–451 (2017).

13. Caravagna, G. et al. Subclonal reconstruction of tumors by using machine learning and population genetics. Nat. Genet. 52, 898–907 (2020).

14. Eirew, P. et al. Dynamics of genomic clones in breast cancer patient xenografts at single-cell resolution. Nature 518, 422–426 (2015).

15. Miles, L. A. et al. Single-cell mutation analysis of clonal evolution in myeloid malignancies. Nature 587, 477–482 (2020).

16. Minussi, D. C. et al. Breast tumours maintain a reservoir of subclonal diversity during expansion. Nature 1–7 (2021).

17. Laks, E. et al. Clonal Decomposition and DNA Replication States Defined by Scaled Single-Cell Genome Sequencing. Cell 179, 1207–1221.e22 (2019).

18. Funnell, T. et al. Single-cell genomic variation induced by mutational processes in cancer. Nature (2022) doi:10.1038/s41586-022-05249-0.

19. Wang, Y. K. et al. Genomic consequences of aberrant DNA repair mechanisms stratify ovarian cancer histotypes. Nat. Genet. 49, 856–865 (2017).

20. Funnell, T. et al. Integrated structural variation and point mutation signatures in cancer genomes using correlated topic models. PLoS Comput. Biol. 15, e1006799 (2019).

21. Li, Y. et al. Patterns of somatic structural variation in human cancer genomes. Nature 578, 112–121 (2020).

22. McPherson, A. W. et al. Ongoing genome doubling promotes evolvability and immune dysregulation in ovarian cancer. bioRxiv 2024.07.11.602772 (2024) doi:10.1101/2024.07.11.602772.

23. Vázquez-García, I. et al. Ovarian cancer mutational processes drive site-specific immune evasion. Nature 612, 778–786 (2022).

24. Kaufmann, T. L. et al. MEDICC2: whole-genome doubling aware copy-number phylogenies for cancer evolution. Genome Biol. 23, 241 (2022).

25. Rose Brannon, A., et al. Enhanced specificity of clinical high-sensitivity tumor mutation profiling in cell-free DNA via paired normal sequencing using MSK-ACCESS. Nat. Commun. 12, 3770 (2021).

26. Ahmed, A. A. et al. Driver mutations in TP53 are ubiquitous in high grade serous carcinoma of the ovary. J. Pathol. 221, 49–56 (2010).

27. Cortés-Ciriano, I. et al. Comprehensive analysis of chromothripsis in 2,658 human cancers using whole-genome sequencing. Nat. Genet. 52, 331–341 (2020).

28. Umbreit, N. T. et al. Mechanisms generating cancer genome complexity from a single cell division error. Science 368, (2020).

29. Hadi, K. et al. Distinct Classes of Complex Structural Variation Uncovered across Thousands of Cancer Genome Graphs. Cell 183, 197–210.e32 (2020).

30. Shen, M. M. Chromoplexy: a new category of complex rearrangements in the cancer genome. Cancer Cell 23, 567–569 (2013).

31. Nguyen, B. et al. Genomic characterization of metastatic patterns from prospective clinical sequencing of 25,000 patients. Cell 185, 563–575.e11 (2022).

32. MacAulay Vacheresse, G., Sabri, E., Domingo, S. & Le, T. Response to subsequent platinum-based chemotherapy post PARP inhibitor in recurrent epithelial ovarian cancer. J. Clin. Orthod. 41, 5578–5578 (2023).

33. Kristeleit, R. S. & Moore, K. N. Life after SOLO-2: is olaparib really inducing platinum resistance in BRCA-mutated (BRCAm), PARP inhibitor (PARPi)-resistant, recurrent ovarian cancer? Ann. Oncol. 33, 989–991 (2022).

34. Patch, A.-M. et al. Whole-genome characterization of chemoresistant ovarian cancer. Nature 521, 489–494 (2015).

35. Stronach, E. A. et al. Biomarker Assessment of HR Deficiency, Tumor BRCA1/2 Mutations, and CCNE1 Copy Number in Ovarian Cancer: Associations with Clinical Outcome Following Platinum Monotherapy. Mol. Cancer Res. 16, 1103–1111 (2018).

36. Cheng, K. W. et al. The RAB25 small GTPase determines aggressiveness of ovarian and breast cancers. Nat. Med. 10, 1251–1256 (2004).

37. Shi, H. et al. Allele-specific transcriptional effects of subclonal copy number alterations enable genotype-phenotype mapping in cancer cells. Nat. Commun. 15, 2482 (2024).

38. Brabletz, T., Kalluri, R., Nieto, M. A. & Weinberg, R. A. EMT in cancer. Nat. Rev. Cancer 18, 128–134 (2018).

39. França, G. S. et al. Cellular adaptation to cancer therapy along a resistance continuum. Nature 1–8 (2024).

40. Wright, S. The Distribution of Gene Frequencies in Populations. Proc. Natl. Acad. Sci. U. S. A. 23, 307–320 (1937).

41. Salehi, S. et al. Clonal fitness inferred from time-series modelling of single-cell cancer genomes. Nature 595, 585–590 (2021).

42. Smith, P. et al. The copy number and mutational landscape of recurrent ovarian high-grade serous carcinoma. Nat. Commun. 14, 4387 (2023).

43. Fischer, A., Vázquez-García, I. & Mustonen, V. The value of monitoring to control evolving populations. Proc. Natl. Acad. Sci. U. S. A. 112, 1007–1012 (2015).

44. Hockings, H. et al. Adaptive therapy achieves long-term control of chemotherapy resistance in high grade ovarian cancer. bioRxiv 2023.07.21.549688 (2023) doi:10.1101/2023.07.21.549688.

45. Gatenby, R. A. & Brown, J. S. Integrating evolutionary dynamics into cancer therapy. Nat. Rev. Clin. Oncol. 17, 675–686 (2020).

46. Zehir, A., Benayed, R., Shah, R. H., Syed, A. & Middha, S. Mutational landscape of metastatic cancer revealed from prospective clinical sequencing of 10,000 patients. Nat. Med. (2017).

47. Huang, X. & Huang, Y. Cellsnp-lite: an efficient tool for genotyping single cells. Bioinformatics (2021) doi:10.1093/bioinformatics/btab358.

48. Layer, R. M., Chiang, C., Quinlan, A. R. & Hall, I. M. LUMPY: a probabilistic framework for structural variant discovery. Genome Biol. 15, R84 (2014).

49. McPherson, A., Shah, S. & Sahinalp, S. C. deStruct: accurate rearrangement detection using breakpoint specific realignment. bioRxiv (2017).

50. Chiang, C. et al. SpeedSeq: ultra-fast personal genome analysis and interpretation. Nat. Methods 12, 966–968 (2015).

51. Weigelt, B. et al. Diverse BRCA1 and BRCA2 Reversion Mutations in Circulating Cell-Free DNA of Therapy-Resistant Breast or Ovarian Cancer. Clin. Cancer Res. 23, 6708–6720 (2017).

52. Lin, K. K. et al. BRCA Reversion Mutations in Circulating Tumor DNA Predict Primary and Acquired Resistance to the PARP Inhibitor Rucaparib in High-Grade Ovarian Carcinoma. Cancer Discov. 9, 210–219 (2019).

53. Li, H. Aligning sequence reads, clone sequences and assembly contigs with BWA-MEM. arXiv [q-bio.GN] (2013).

54. Scheinin, I. et al. DNA copy number analysis of fresh and formalin-fixed specimens by shallow whole-genome sequencing with identification and exclusion of problematic regions in the genome assembly. Genome Res. 24, 2022–2032 (2014).

55. Schubert, M. et al. Perturbation-response genes reveal signaling footprints in cancer gene expression. Nat. Commun. 9, 20 (2018).

56. Medina-Martínez, J. S. et al. Isabl Platform, a digital biobank for processing multimodal patient data. BMC Bioinformatics 21, 549 (2020).

